# An incremental training method with automated, extendible T-maze for training spatial behavioral tasks in rodents

**DOI:** 10.1101/514703

**Authors:** Esther Holleman, Jan Maka, Tim Schröder, Francesco Battaglia

## Abstract

We present a training procedure and a T-maze equipped with sensors and automated feeders for training spatial behavioral tasks in rodents. The maze can be transformed from an enclosed box to a maze of variable dimensions. The modularity of the protocol and setup makes it highly flexible and suitable for training a wide variety of spatial tasks, and facilitates incremental training stages of increasing maze size for more efficient learning. The apparatus, in its software and hardware, is able to adapt to animal performance, adjusting task challenges and difficulty.

Two different methods of automatic behavioral scoring are evaluated against manual methods. Sensors embedded in the maze provide information regarding the order of reward locations visited and the time between the activation of the cue via the nose-poke and the activation of the reward location sensors. The distributions of these reaction times differ between correct and incorrect trials, providing an index of behavior and motivation. The automated maze system allows the trainer to operate and monitor the task away from the experimental set-up, minimizing human interference and improving the reproducibility of the experiment. We show that our method succeeds in training a binary forced-choice task in rats.

## Introduction

Spatial behaviors are particularly developed in rodents as they have an innate drive to explore new environments and navigate through narrow passageways [1, 2]. For this reason, many cognitive functions, including those that are not necessarily spatial in nature, such as working memory, are examined using tasks with a strong spatial component [3]. Place cells in the hippocampus are a useful model in the study of memory [4]. These neurons, among others, have spatially tuned responses, providing a neural read-out of behavior. Accurately assessing the spatial tuning of those cells requires large mazes, typically 80-100 cm or larger, rather than smaller, Skinner-box like setups, most convenient for e.g. operant conditioning training. The size of the maze does not present a problem for simple, exploratory tasks. However, shaping rodents into producing specific behavioral responses to cues can prove challenging in large environments that entice rodents to explore.

With setups typically used for behavioral electrophysiology, training these tasks is often labor-intensive, requiring the trainer to be near the maze to give cues, open and close barriers, and the reward the rat when it displays the desired behavior [5-7]. Furthermore, experimenter proximity and involvement can influence the performance of the task. The trainer may unintentionally signal the correct answer to the animal, as a result the animal may be responding to unintended cues of the trainer instead of the intended cue [8]. This is known as the experimenter-expectancy effect, and was first documented by Oskar Pfungst in 1907 in the case of the horse known as ‘Clever Hans’ [9]. These possible confounds can make it more difficult and time intensive to train a task, and reduce the reproducibility of the task. These unwanted effects can be largely resolved through the use of automated training systems such as B.F. Skinner’s operant conditioning chamber [10]. These systems have evolved for various applications, and can include touch screens [11], acoustically transparent chambers [12] and high throughput systems implemented in the home cage [13, 14]. The various custom made and commercially available automated training systems available are mostly implemented in Skinner boxes however. Automated systems designed for tasks in larger mazes are less commonly seen.

However, designing an automated system for a large maze is subject to different constraints than those posed by a Skinner box. The enclosed environment of a Skinner box provides an environment with minimal distractions. A similar effect could be achieved in a larger environment by implementing high, opaque walls around the track. This is not a possible solution for all tasks however, as the visibility of spatial cues is paramount for the formation of spatial representations, e.g. in the hippocampus. Another advantage of smaller training spaces such as Skinner’s operant chamber is that the cue can be immediately followed by a reward, facilitating association. In contrast, the likelihood of distracting events taking place between the cue-delivery and the discovery of the associated reward is increased significantly in a larger maze, because of the longer trial duration and the larger number of environmental stimuli, making it more difficult for it to make the association between the cue and the reward. In a large maze, these additional difficulties are exacerbated when all task components are trained simultaneously from the start of training. A task such as visiting a reward area associated with a particular cue tone consists of several subtasks that the animal must acquire. First the animal must learn to nose-poke to initiate a trial and receive the cue, learn that only one of the two reward areas can be visited in order to obtain a reward, and learn to return to the start-box and nose-poke to initiate the next trial, regardless of whether reward was received in the current trial. Attempting to train all these aspects at once in a large maze, where the animal can easily be distracted and the cue tone long forgotten before the distant reward area is reached, needlessly increases the difficulty of the training process.

We propose a method to facilitate training complex tasks in a large environment by dividing the training into stages of increasing difficulty, and have created an automated setup that streamlines this procedure. Early training stages should minimize distractions and minimize the time between cue and reward to facilitate the forming of an association. The distance between the cue and reward can gradually be increased in each training stage until the final task to be trained is reached. Thus, in early training stages the reward location should be located close to the start-box where the cue is received to encourage the development of an association between the two. When the response of the animal to the cue indicates the associations have been formed the distance between the cue and reward locations can incrementally be increased to increase the difficulty level of the task and the working memory load. The initial close proximity of the cue and reward locations also enable the creation of a more sheltered environment, decreasing the amount of possible distractions during early training stages, facilitating efficient learning.

Applying this gradual training schedule requires adapting the physical maze and the training software as the animal learns. To implement this method, we have created an automated training system consisting of a modular maze, hardware, and a software program to run training sessions. The modular nature of the maze facilitates incremental training stages of increasing maze size. At the beginning of each block of trials the software generates a randomized, counterbalanced trial schedule. Timing and delivery of cues is controlled by a micro-controller, receiving information from sensors located at relevant locations in the maze, and running each trial autonomously, with no involvement of the controlling computer It is possible to override the decisions made by the micro-controller algorithm through the user interface of the software. Feeder units can be activated and trials can be canceled in this manner. This ability to intervene from a distance eliminates the need for the researcher to physically enter the experiment room and risk affecting the performance of the animal [15,16]. This system has been used here for a spatial working memory task on a T-maze, however it can be implemented on various maze designs for a variety of tasks. Possible modifications include adding a visual cue, visual feedback, or positioning two feeder units by the nose poke to provide cues in the form of flavored pellets and providing pellets of a flavor corresponding to the cue at the reward areas. The amount of reward areas can be increased through the addition of feeder units, for instance in the case of a radial maze with several arms. The system easily accommodates for the flexible placement of multi-modal cues and for different cue-reward schedules, therefore lending itself to a number of different experiments. We demonstrate its functionality here by training a tone-to-place association task (see e.g. [6]).

## Methods

### Animals and behavioral task

All animal procedures were approved by the Animal Ethics Committee of the Radboud University Nijmegen (RUDEC) and carried out in accordance to the Dutch guidelines and regulations for animal experiments.

Four six months old Long Evans male rats were housed in pairs and maintained on a reversed 24h light/dark cycle at 85% of their ad libitum weight and trained on a two-alternative forced choice task in a T-maze to test working memory (see p18 supplementary materials on animals and food restriction for details). Upon activation of the nose poke by the animal either a 7Hz or 14Hz tone was played as a cue to indicate which location would be rewarded when visited.

In the early phases of training the distance between the nose poke and the reward areas was minimized to facilitate the formation of an association between the tones and reward areas. This distance was increased incrementally according to the performance of the animal, with the maze gradually morphing from an operant conditioning box into a full-blown T-maze (Figure 1).

**Figure 1.**
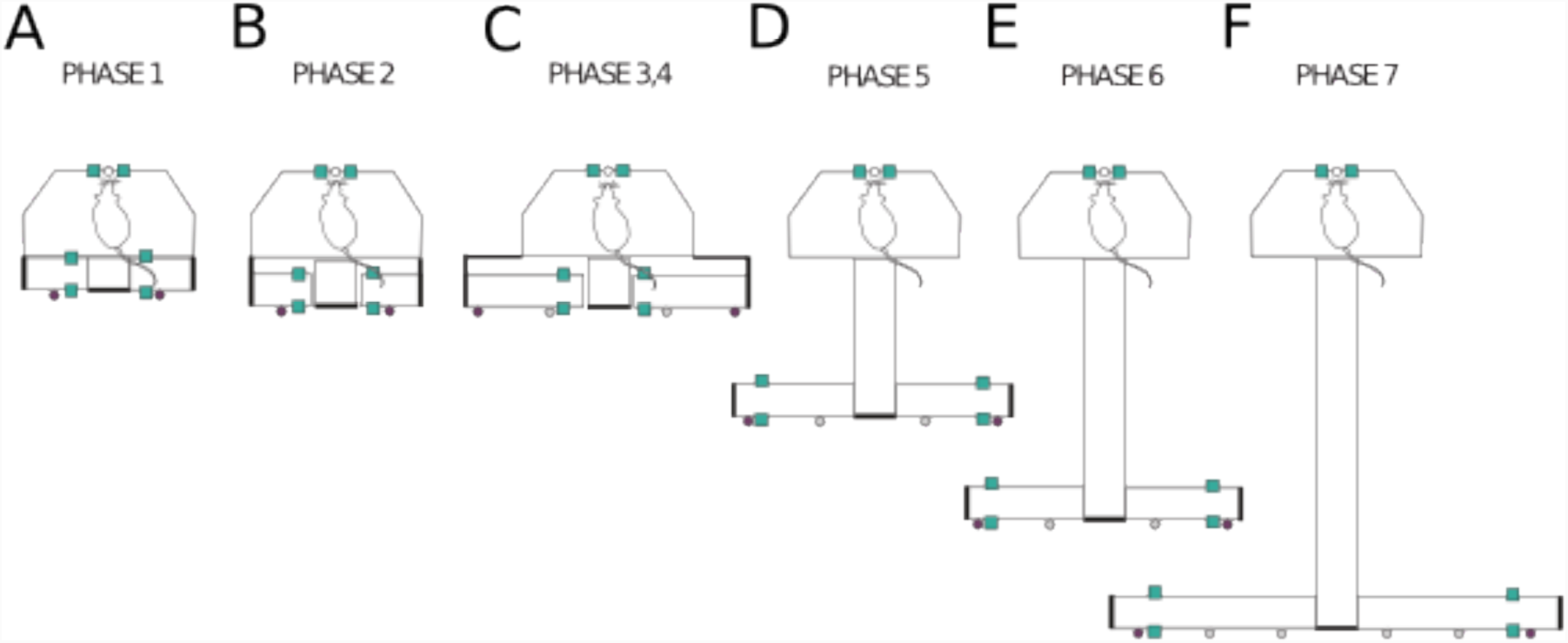
Configuration of training environment at each phase. Locations of infrared sensor are shown as teal squares at the nose poke and the reward areas. Feeder locations are indicated by filled circles (empty circles represent possible feeder locations). **A)** In phase 1, a small, enclosed space is created ensuring the cue and reward are administered in close proximity. **B)** A gap between the arms of the maze and the start box platform is introduced in phase 2. **C)** The side arms increase in length at phase 3. **D)** At phase 5 the central arm is increased in length by 20 cm, the length of the sidearms is increased to 20 cm and the reward areas, including sensor locations shift to the end of the side arms. **E)** At phase 6, the length of the central arm is increased to 50 cm and at phase 7 to 80 cm **F)** Also at this phase, the arms reach their maximal length of 40 cm and the distance between the central arm and the reward areas is increased further to 35 cm.

### Training protocol

Training began with habituation to the maze, followed by nose-poke training, automated feeder habituation, cue training, and ultimately the performance of the complete task in increasing levels of difficulty. To enable gradual acquisition of the different task components, training was divided into seven phases (Table 1; see Supplementary material for an extended description of the training phases). The first four phases consisted of cue training, followed by three phases where the animals performed the complete task that included a working memory component facilitated by increased distance between the nose poke and reward areas. Each training session was divided into four blocks. The number of trials per block increased from 10 trials during the first phase, to 15 trials for phases two through five, 20 trials during phase six, and 25 trials in phase seven.

**Table 1.** Training phases

| Overview of Training Phases |  |
| --- | --- |
| Phase | Learning Goal / Desired Behavior |
| 1 | The task structure is acquired. Initialize trial through nose-poke, visit one of two reward areas, consume reward, and return to nose-poke for subsequent trial. Hint trials, where a pellet is dispensed at the reward areas associated with the tone, are used initially to encourage the animal to visit the reward area following a nose-poke. The time between the |
|  | tone and the hint is gradually increased. When the animal displays any movement towards the correct area before the hint has been given three pellets are rewarded at that area. |
| 2 | Respond to the cue tone through head movement in the direction of the associated reward area. |
| 3 | Respond to the cue tone with complete body turn towards associated reward area. |
| 4 | Respond to the cue tone with complete body turn and forward movement towards associated reward area. |
| 5 | Maze arms moved back 20cm from start-box. From this phase onwards the animal must respond to the cue tone by moving towards one reward area through the central arm and waiting at reward area for the reward. |
| 6 | Maze arms moved back 50cm from start-box. |
| 7 | Maze arms moved back 80cm from start-box. |
| **Table 1:** Training phases |  |

### Randomization for sequence of rewarded areas within a block of trials

A sequence of random integers (0 or 1) was generated to determine which side to cue/reward in each trial. Sequences with consecutive repetition of a particular side for more than 3 trials were discarded to prevent the animals from developing a bias towards a particular side. Similarly, sequences with frequent switches of rewarded sides between trials were also removed to reduce the natural tendency of the animal to alternate between reward sites with each trial [17]. This alternation strategy commonly applied by rodents may reflect natural foraging behavior where it is not strategic for the animal to return to a depleted food source [18]. See supplementary material for a full description of the algorithm.

The randomization algorithm was tested by generating the sequence of cues for a block of trials ten thousand times. For each generated sequence several measures were tested such as the percentage of left versus right rewarded trials, the amount of alternation between sides, the ratio of left to right transitions, and the ratio of right to left transitions (Supplementary Methods p8-10, Supplementary Fig. 6). If any patterns were found that could potentially encourage a bias the algorithm was altered until no potential biases were present. We found, however, that a ratio of 40% alternation transitions to 60% same side transitions was most conducive to learning as the alternation tendency of the rodents was sufficiently strong that it must be actively discouraged by presenting the animal with more same-side transitions than alternation transitions. To study spatial memory in animals it is imperative to ensure that the behavioral readouts can be attributed to spatial associations to the cue and are not the result of other underlying strategies that produce similar behavioral readouts [19]. For example, rats are excellent in identifying patterns in the randomization sequence in order to predict which side has a high probability of delivering reward in the upcoming trial, given the outcomes of the previous trials [20]. To ensure animals could not acquire too many rewards by using a strategy other than the task to be learned, simulations using various strategies were performed on the sequences of trials generated with the randomization algorithm. Both simple strategies such as consistently choosing one side over the other or spontaneous alternation between reward areas as more responsive strategies such as selecting the reward location opposite to the one rewarded in the previous trial (win-shift) or returning the location rewarded in the previous trial (win-stay) were tested. The sequence generator was deemed sufficient when the strategy simulations could not score above 60% on average over any of the session lengths used in the experiment.

### Analysis methods

For scoring method analyses shown in figure 3, outliers were removed by first performing a linear regression and calculating the standard deviation of the absolute value of the distances between the data points and the regression fit. Data points located further than 3 times the standard deviation away from the regression fit were marked as outliers. Subsequently, another regression fit was performed through the remaining points. This second fit is shown in the subplots, outliers are displayed as open circles.

Timed-out trials were scored as incorrect, however they were not included in the reaction time density distributions as they are all set at the predetermined threshold, for example at six seconds, and therefore not representative of actual animal behavior. Also, due to the inaccurate sensor readings in earlier phases only reaction times from the full task, phases five to seven were included.

Learning was assessed through automatically calculated scores. These scores were compared to manual scores throughout training to ensure accuracy. To determine if correct scores were due to animals behaving according to the task or according to a strategy, animal scores were tested for compliance with common strategies such as win-shift, win-stay, and spontaneous alternation strategies.

More detailed accounts can be found in the Supplementary Methods for the training protocols (p1-4, Supplementary Table 1, Supplementary Figure 1), the randomization testing (p8-10, Supplementary Figure 6), strategy simulations (p10-13, Supplementary Figures 7 and 8) and compliance of animal responses to strategy responses (p14-15).

## Results

The learning curve of the animals over the different phases is shown in figure 2A. The percentage of correct trials is shown on the y-axis and the training days on the x-axis, where the phase in which the training day occurred is also displayed. An increase in performance over phases can be observed. The average score across animals increased gradually from 59% correct in the first phase to 87% correct in the last phase.

**Figure 2.**
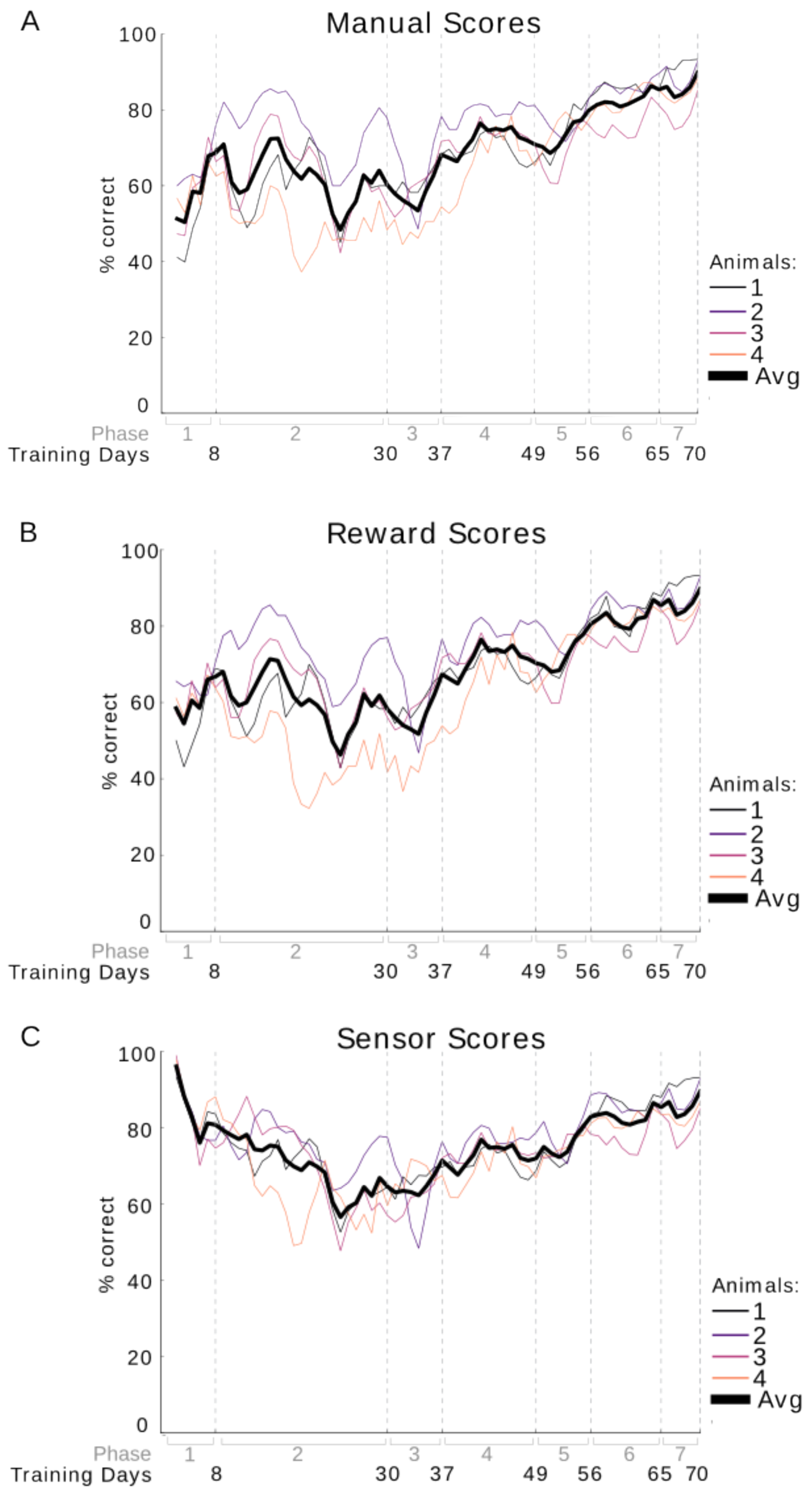
Comparison of scores calculated manually during training, scores acquired by counting the number of pellets rewarded, and scores as recorded by the sensors. A rolling average over 3 days was used to calculate the percent of correct scores over the training days (training phase shown in gray). The scores for each animal are plotted separately and also as an average over all animals (thick black line). Scores based purely from sensor readings are unreliable for the first four phases as the tail of the rat could activate the sensors unintentionally. Additionally on the first two days of the first phase reward automatically followed the cue in many trials. These hint trials were excluded from the scoring process however they do occur in the sensor data, resulting in unusually high sensor scores in the first phase. **A)** Results from manual scoring. **B)** Scoring based on amount of rewards given to the animal. In phase 1, the criteria for a correct trial was set at three or more rewarded pellets, in phase 2 to 5, two or more pellets, and in phase 6 and 7 one or more pellets. **C)** Scores based on sensor data from sensors at the reward locations. Activation of a sensor at the reward area corresponding to the cue resulted in a correct trial.

**Figure 3.**
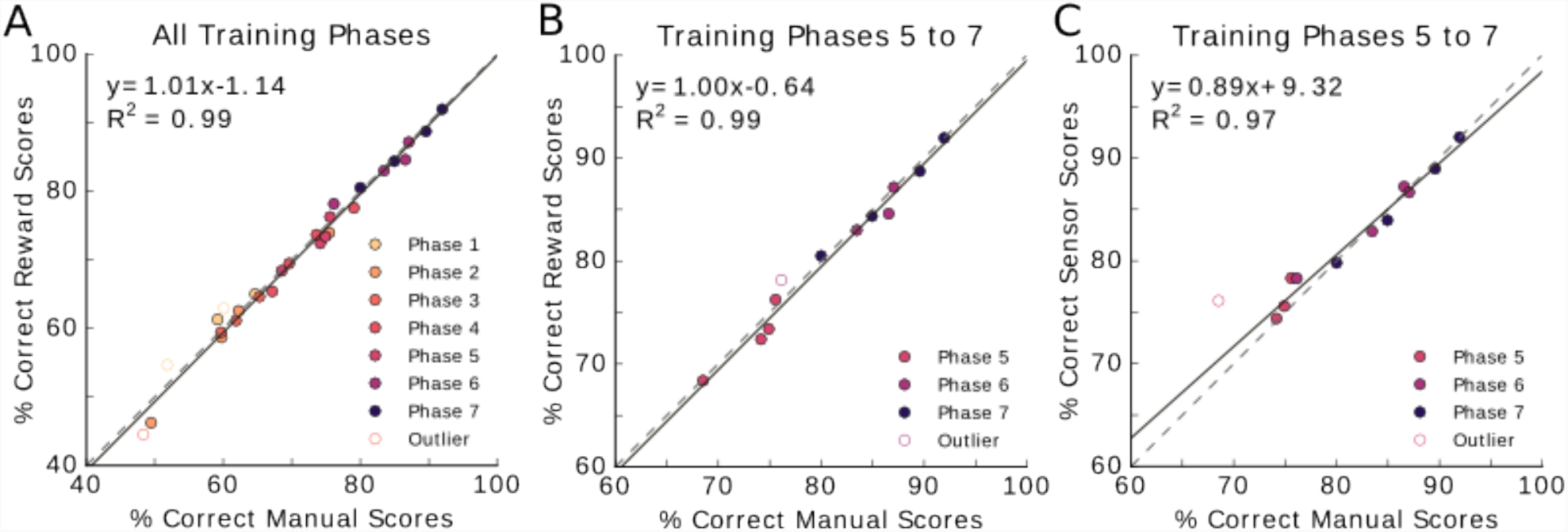
Scoring methods compared in scatter plot. Each phase contains four data points, representing the average score obtained during the specified phase for each of the four animals. Phases are depicted in different colors. Outliers, indicated by open circles, were removed for the regression analysis (see Methods). The dashed diagonal line indicates the unity line, where both scores are identical. The solid line represents the linear regression fit through the data points. **A)** The percent correct for manual scores are plotted against the percent correct for the reward-based scores. All data points lie close to or on the unity line indicating the similarity of these scoring methods (slope = 1.01, y-intercept = -1.14%, R^2^ = 0.99, p=2.03×10^-25^). **B)** Reward scores as a function of manual scores (slope = 1.00, y-intercept = -0.64%, R^2^ = 0.99, p=1.26×10^-9^). **C)** Sensor scores as a function of manual scores (slope = 0.89, y-intercept= 9.32%, R^2^ = 0.97, p=1.82×10^-8^.

The learning curve in figure 2A was acquired using manual scoring to mark a trial as correct or incorrect.

### Comparison of scoring methods

To test the accuracy of the automated scoring, scores were manually recorded during training (Figure 2A). Two methods of automated scoring were compared, scoring based on the number of rewards given (Figure 2B) and scoring based on sensor information (Figure 2C). As described previously, training this task involved rewarding based on performance. The first and second phase of the cue training included ‘hint’ trials where one reward was given immediately following the cue tone. In the manual scoring, hint trials were excluded. However, as the animal often did move to the correct reward area following a hint, the sensors record these trials as correct. The reward scoring does not count hint trials as correct, as only one pellet is delivered during these trials compared to several pellets in trials where the animal chose the correct location without the help of a hint. With the reward scoring method a trial was marked as correct depending on the amount of reward given. The amount of rewards necessary to qualify a trial as correct depended on the training phase. In phase 1, the threshold for a trial to be considered correct was set at three or more rewarded pellets. In phase 2 to 5, two or more pellets, and in phase 6 and 7 one or more pellets.

The similarity between the learning curves of the manual scoring method and of the reward scoring method (Figure 2A and B) indicates that reward scoring is a suitable replacement for manual scoring. Linear regression analysis also supports this, as the fit has a slope of 1.01, indicating the results for manual and sensor scoring methods are close to identical (Figure 3A, R^2^ = 0.99, offset -1.14%, p=2.03×10^-25^).

From phase five onwards, both manual and reward scoring methods (Figure 3B, R^2^ = 0.99, p=1.26×10^-9^) as well as manual and sensor scoring methods are highly correlated (Figure 3C, R^2^ = 0.97, p=1.82×10^-8^). In both cases, the data scattered around the unity line with slopes of 1.00 and 0.89 respectively, and offsets of -0.64% and 9.32%, respectively.

### Animal behavioral responses in light of common strategies

Animal responses to the cue were compared to strategies commonly applied by rodents in binary choice tasks to ensure animals learned the task intended to be taught. Given the response of the animal to the first cue in a block of subsequent trials the response for the remaining trials was predicted for several common strategies. These strategy responses were then compared to the actual responses of the animal.

### Reaction time distributions

A trial starts when the nose-poke sensors are activated and ends either when the rat crosses the sensors at one of the reward areas. A trial is also terminated when the maximum trial time is reached before reward sensors are activated. The thresholds for this time-out differ per phase. In the first phase, the animal has six seconds to make his way to the reward areas, as the difficulty increases with each phase the time allowed per trial decreases. The distributions in Figure 5 show the reaction times for each animal for both correct and incorrect trials (left to right) recorded in phases five through seven (top to bottom). The number of trials included in each distribution is noted per rat.

**Figure 4.**
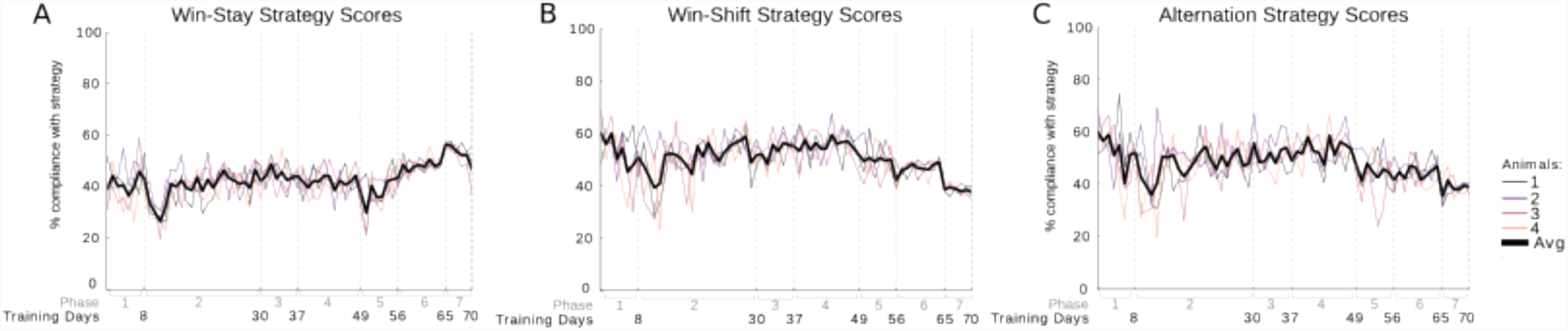
Animal responses compared to strategy responses. Percent compliance to strategy (y-axis) over training days (x-axis, phases indicated in grey). **A)** Responses for each animal compared to expected responses when applying a win-stay strategy. **B)** Animal responses compared to win-shift strategy responses. **C)** Animal responses compared responses expected for spontaneous alternation.

**Figure 5.**
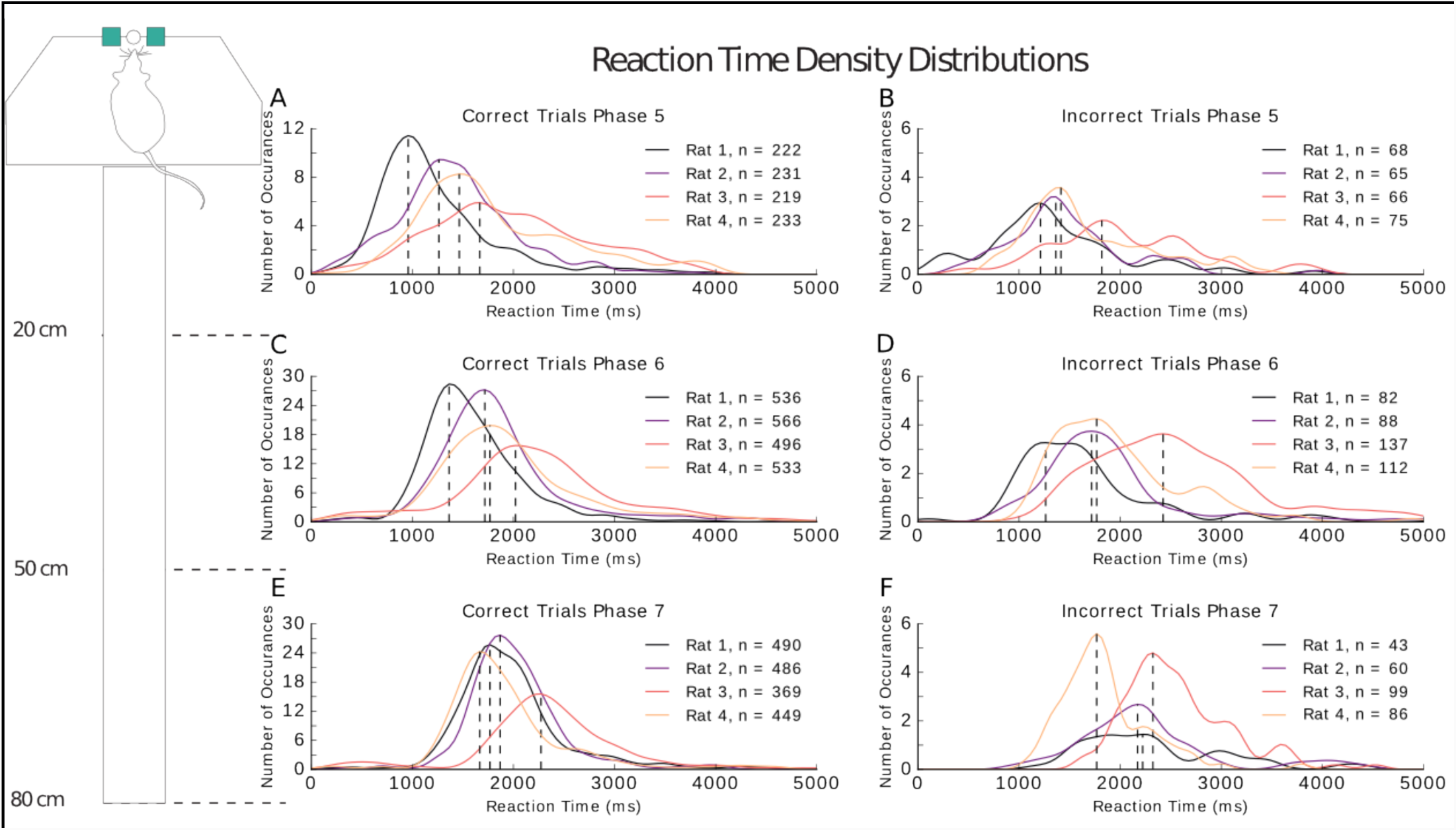
Reaction time density distributions for the full task (phases 5 to 7). Distributions are plotted per rat. The number of trials included in each distribution are noted in the legend. Reaction time is shown in milliseconds. Correct trials are plotted in the right column and incorrect trials in the left column. Each row represents a phase. The dotted lines are the modes for each distribution.

From the shape of the distribution information about the behavior of the animal may be inferred. A broad curve with widely varying reaction times indicates inconsistent behavior. Slow reaction times were often due either to hesitation following the administration of the cue or distracted behavior where the animal was clearly disinterested in the task and likely no longer motivated by the food reward. In those trials, animals only performed at chance level. An unusually fast response points towards the animal following a non-task-related strategy, for instance the animal may have based its choice on the result of the previous trial and therefore had no need to attend to the cue. The reaction time distribution of all incorrect trials spans over a large range of values, as it covers the fast reaction times, likely reflecting the use of a strategy, to the slow reaction times presumably related to indecision, virtual trial-and-error behavior [21], or lack of motivation. In comparison, distributions of correct trials often cover a much smaller range of reaction times, resulting in a narrower distribution. Reaction times of animals that have successfully made the association between the cue and the reward areas vary between 1000 and 2000 milliseconds. This average increases with the size of the maze, as the distance between the nose-poke and the reward areas increases.

The differences between the correct and incorrect reaction times increase as the phase of training progresses (phase 5: p=0.16, phase 6: p=0.0001, phase 7: p= 7.2 x 10^-9^ two-sample Kolmogorov-Smirnov test). In Figure 5A the reaction times of all trials in phases 5 through 7 show a significant difference between incorrect (green) and correct (blue) trials (p=1.24 x 10^-7^ two-sample Kolmogorov-Smirnov test, interquartile range correct trials = 558 ms, incorrect trials = 890 ms). Figure 6B shows that the distributions for the correct trials narrow with each successive phase. The interquartile range for correct trials is 840 ms in phase 5, 650 ms in phase 6, and 520ms in phase 7. For the incorrect trials this effect is not present (phase 5: 765 ms, phase 6: 746 ms, phase 7: 669 ms).

**Figure 6.**
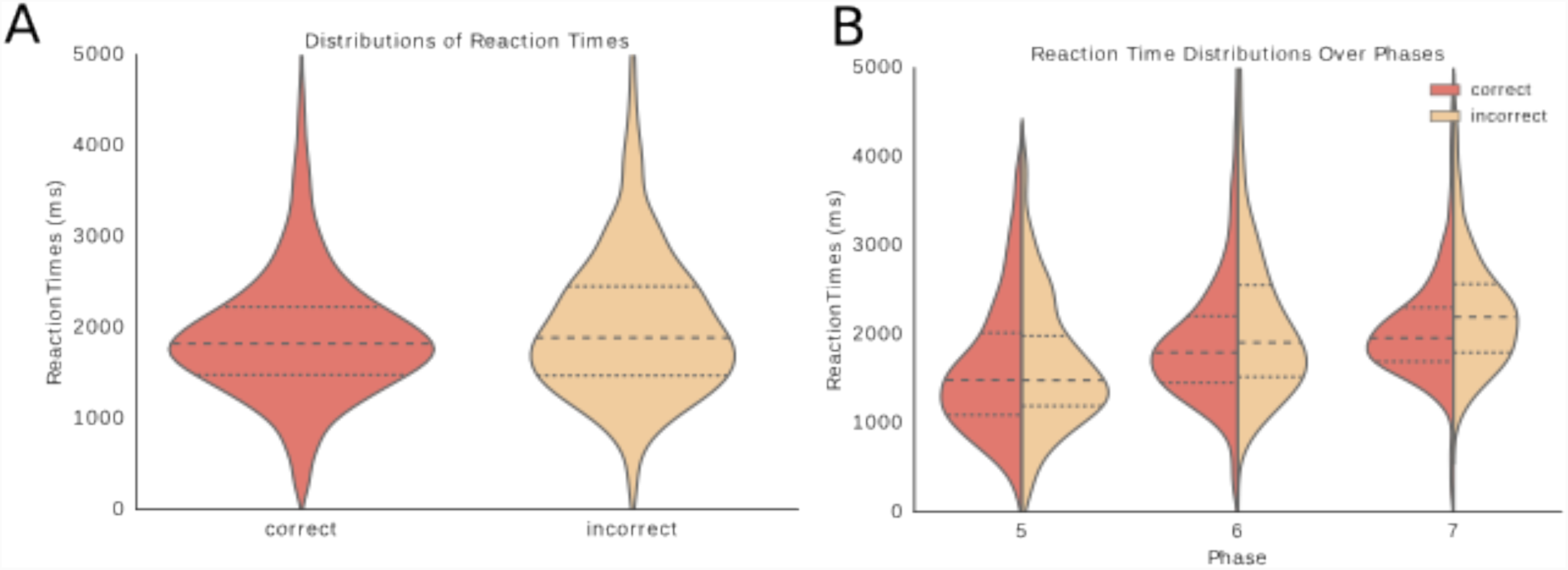
A) Distributions of reaction times in milliseconds for phases 5 to 7 plotted together to compare correct trials to incorrect trials (p=1.24 x 10^-7^, two-sample Kolmogorov-Smirnov test). **B)** Reaction time distributions for each phase separately with reaction times in milliseconds plotted on the y-axis and phases on the x-axis. The left half of the violin plots represent the reaction times of the correct trials and the right side those of the incorrect trials. Reaction times increase over phases as the distance travelled increases with the expansion of the central arm of the maze.

## Discussion

We have presented a novel training procedure and setup for maze-based tasks, which enables the modular extension of the training apparatus, facilitating learning. The setup is automated, with the within-trial logic controlled by a micro-controller, with the overall experimental logic overseen by a PC, enabling user intervention when needed. Micro-controller based automated mazes and training setups have recently been proposed [22-24]. Here we propose a modular approach (with “slave” micro-controllers handling sensors in each maze component and a “master” component coordinating the trial management), enabling the easy extension and customization of the environment. All components are low-cost and readily available, and assemblage of the system is easy.

We have shown that rats can learn the sound-to-place association task when trained through this system. The first phases, which consisted of learning the associations between the cues and the reward areas were the most time consuming, and those took place in the reduced-size version of the setup. The gap created when the arms are moved away from the start box only appeared to affect the behavior of the animals when first introduced in the third phase. Once the animals were accustomed to the gap between the maze arms and the start-box moving the arms further away incrementally did not affect their performance. Furthermore, none of the distances used (up to 80 cm) appeared problematic for working memory. Rats were able to complete 100 trials a day with ease when number of trials was increased gradually during training. Notably, this was based on appetitive food rewards, whereas other approaches to obtaining large number of training trials involved invasive stimulation of reward structures (e.g. the medial forebrain bundle) for reward [25]. Choosing a task that facilitates a large number of trials per day allows for the acquisition of a sufficient amount of behavioral data for statistical analysis and is beneficial for electrophysiological data acquisition. The automated training system can be implemented in many different tasks and mazes. It is also possible to work without a cue and simple give reward at a particular area in the maze in which the sensors have been activated. The amount of different cues and reward areas can be extended through the addition of more feeder and sensor units. Variations of cues and rewards are also possible. For instance, feeders can be attached to the nose poke device to feed different flavors of pellets as cues instead of the cue tones used in this experiment. The feeders at each reward area can also contain different flavors to match the cue flavor. Possible extensions of the setup include the addition of an odor distributor to provide cues, or a display to provide visual cues.

Bower and McNaughton show that training the same task with different methods resulted in differential hippocampal encoding [5]. Rats were trained to remember sequences of reward locations containing repeated locations within different sequences in a delayed alternation task on a continuous T-maze. For instance, location ‘b’ (the central arm of the T-maze) is present in both the sequences ‘“a-b-c” and “d-b-e”. The rat must differentiate between the two contexts in order to decide which location to visit after location ‘b’. When animals were trained, as an intermediate step, by also placing reward at the end of the repeated segment, they learned the sequences correctly, yet hippocampal ensemble activity did not differentiate the sequential context of the repeated segment. In rats that learned the task with the help of moveable barriers that directed their path to the reward areas hippocampal activity did differentiate between the different sequential contexts. The method proposed here increases maze size and trial duration, while leaving the structure of the task unchanged. We think that this may facilitate the study of neural representations of rule learning and task structure.

An automated training system lends itself well to examining the effects of various training methods on neuronal activity as it offers a controlled way to train these different methods, without interference. For example, the same working memory task described here could be trained with and without barriers placed in the maze, directing the animals to the correct reward location. Similarly, comparisons between various forms of cueing could be examined, such as cue LEDs located at the reward areas, or cue LEDs on either side of the nose-poke device, indicating which side will be rewarded. This requires a system where a task can be trained both with and without barriers such as automated doors. However, most automated mazes employ doors to enforce trial structure, for example to restrain the animal in the start-box before the initiation of a trial, to ensure it does not visit multiple reward areas after an incorrect behavioral response, and to direct the animal back to the start-box [26, 14]. We show here that it is possible to train rats to both conform to the trial structure and perform the desired task through incremental training phases without barriers, thereby facilitating experiments where the use of barriers may influence the neural processes to be studied, yet automation of the trial structure is desired.

Several measures can be taken to improve the experimental set up described in this paper. These include creating more consistency among phases by introducing a gap between the start-box and the maze arms during habituation and increasing the amount of training sessions to two per day in all phases to expedite the learning process and improve the accuracy of the analysis. Although block length should remain short in the first phase to provide enough time for the animals to habituate to their new routine in a stress-free manner, increasing the amount of blocks of trials per day could significantly accelerate the learning process and decrease the amount of training days necessary, provided enough breaks are present between blocks. In this experiment, rats were trained in half-day sessions for the first five phases. Experience in later stages showed no loss of focus, motivation, or performance in the animals when two training sessions were provided daily, one morning and one afternoon session, with a break of at least two hours in between, allowing the animals to rest, drink, and regain motivation. Another benefit of increasing the amount of trials per day is that it allows for feeding the animals their daily amount of food required to maintain a healthy weight within the task and avoids the necessity to feed extra in the cages after training sessions. This circumvents problems with dominance between animals housed together [27, 28], where the dominant animals consume more food in the cage and perform poorly as a result due to lack of motivation, while the performance of non-dominant animals suffers from anxious behavior during the task as a result of underfeeding. Currently, in the first three phases the maze arms are placed close to the start-box. This, however, encourages the rats to take a shortcut to the reward areas. Observations from this experiment indicate that it would be more beneficial to move the arms a short distance away from the start-box during habituation. This will provide the animals with more time to become acquainted with the gap between the arms and the start-box during the habituation period instead of between training phases, as a result the performance of the rat will be less influenced by transitions between phases. Moving the arms a short distance from the start-box initially will also require the animals to travel via the central arm to arrive at the reward locations for all phases of training instead of only in the later phases. This creates consistency across phases that will ease the interpretation of neural data and facilitate the automation of the entire task, as sensor data is reliable from the first phase onwards, resulting in more reliable analyses of all phases.

In conclusion, we presented a novel framework for animal training in spatial, maze-based task, which may prove itself useful for those attempting to train animals in the large setups that are needed for example for neural ensemble recording experiments.

## Competing Interests

The authors declare no competing interests.

## Author Contributions

EH and FPB designed research, JM, TS and EH developed hardware and software, EH performed research, FPB supervised research, EH and FPB wrote the paper.

## Data availability

All data, software and hardware schematics are available upon request.

## Acknowledgements

We thank Ambra Marelli, Ana Menegollai, and Iris Villani for help with the construction of the maze and the training of the animals during their internships, Dr. Peter Bremen for valuable insights regarding the analysis and Dr. Lisa Genzel for insightful comments on this manuscript. The research leading to these results has received funding from the European Union’s Seventh Framework Programme (<u>FP7/2007–2013</u>) under grant agreement no. 284801 (Enlightenment) and no. 600925 (NeuroSeeker).

## T-maze task – General training description

The desired behavior to be trained in this task involves the presentation of a cue tone, either 7 or 14 kHz, in response to which the rat must move to the location associated with the tone. The 7 kHz tone signals that the rat must move to the left reward location, in the case of a 14 kHz cue it must move to the right reward location. When the rat moves to the correct location pellets will be delivered at the chosen reward location. If the rat chooses incorrectly, no reward will be provided. Also, if the rat chooses the incorrect location first and immediately proceeds to the correct location afterwards it will not be rewarded as the task dictates that the correct location must be visited first. This is implemented through the use of sensors in the maze. The pair of sensors activated first will report their activation to the main program and prevent any other sensor activation from triggering a reward.

At the core of this task lies the association between the cues and the reward areas. The rat must learn that tone played at the beginning of each trial, initiated by a nose poke, is related to receiving reward from a particular reward location. Furthermore, the rat should also move to the reward location in response to the cue in order to receive the reward at that location. A delay between the stimulus presentation and the choice leading to a reward can be used to test working memory.However, efficient initial training for this behavior and successfully requires a short amount of time between the cue and the reward for an association to be made between the two.

The modular structure of the maze with adjustable arms, enables the successive expansion of the maze to larger sizes. These arms contain the reward areas, and can be moved increasingly further away from the nose poke, where the rat receives the cue. Initially the arms are located directly next to the startbox platform, minimizing the distance and therefore time, between the cue and the reward. The startbox and perimeter of the maze are equipped with 40 cm high walls to limit distractions from external stimuli.

The task structure is established during the first training phases. This involves nose-poking and moving to a reward area before nose-poking again for the next trial. Before moving to the next phase, the behavior of the rats should follow the task structure, always nose-poking first, moving to a reward area, consuming pellets at reward and returning to the nose-poke.

Next the association between the cue, the desired behavior, and the reward must be established. Compared to later phases this phase is the most time consuming (29 days, see Table 1 for a comparison with other phases). In order to build a strong association, the rat must actively choose a side in response to the tone in order to receive a reward. In this phase the rat must always first move towards the cued side. At first a reward will follow after only a small movement of the head towards the correct reward location, the required movement to trigger reward will become increasingly larger, leading up to the point where the rat must move to the reward location and wait there before receiving a reward.

Once an average performance of 70% correct is achieved over at least 3 training days the maze configuration can be adjusted by moving the arms further away (see Supplementary Figure 1) from the start box. The rat should then be trained in the same manner, continuing the same task as before in the new configuration, until performance is satisfactory.

The maze arms, and therefore reward locations can be moved increasingly further away gradually, each time ensuring the rat reaches a consistently good performance before moving to the next phase. As a result, the decision point will also be moved increasingly further away from the cue. Whereas at first rats pointed towards one location or the other directly following the cue, they must now walk down a central arm for an increasingly large distance before committing to a choice. In this task configuration, choosing the correct location requires holding the cue flavor in working memory. This will be trained in several phases where the distance for the rat to hold the cue in memory will be increased incrementally with every phase.

Below, we provide a detailed protocol for training:

## 1. Habituation to maze & the association of reward locations with food

1. Thoroughly handle animals, getting them used to the experimenter and to being picked up, placed back into the home cage, until this produces no apparent stress
2. Begin the habituation to the training environment by placing group-housed, food restricted animals in the maze together for a short period of no longer than 2 minutes with pellets available at reward areas and at the edge of the nose poke tube. Following a break of 5 minutes in the home cage the rats can be placed back in the maze and left there for one minute longer than the previous session. Repeat this process until rats can comfortably be in the maze together for 6 minutes. Moving back and forth from home cage to maze will also accustom the animals to the process of entering and exiting the training environment. The goal of these sessions is to generate an association between the training environment and food reward. If at any time an animal displays signs of distress, remove the animals from the maze, provide a 5 minute break, and repeat the previous session until the animals can comfortably remain in the maze for the desired time. Due to a neophobia towards food found in novel environments the animals may not eat the food on the first day [1].
3. Repeat this process the following day, leaving the animals in the maze together for 5 minutes initially, followed by a 5 minute break in the home cage. Repeat this process further while incrementing the time in the maze by one minute per session.
4. The following two days repeat the above, starting with an in-maze time of 2 minutes on the first day, with one animal at time to accustom the animals to spending time in the maze alone for increasingly longer periods of time. On the second day begin with a 5-minute session.
5. Only move an animal on to the nose poke training when it consumes the pellets provided in the training environment. If this is not the case after the fourth day repeat the 10-minute sessions until the animal is comfortable eating in the maze. Growths curves should be carefully monitored and the optimal percentage of normal weight for each animal identified and maintained throughout the duration of the training.

## 2. Nose-poke training

Place pellet near the edge of the nose poke tube. Wait until the rat consumes this pellet. Repeat this several times before placing the pellet slightly farther up the nose poke tube. Gradually increase the distance of the pellet from the entrance of the nose poke tube until the animal is nose poking far enough into the tube to activate the sensors.

Once the rat displays nose-poking behavior correctly in a consistent manner, follow the nose-poke reward immediately with a reward at one of the reward areas. Do not reward in the nose-poke again until the rat has visited the rewarded location. Randomize the locations rewarded in such a way that the rat visits both equally often (see ‘Randomization’ section for details). The rat must always nose-poke before receiving a reward at a reward area. Bait the nose poke for 5 trials, and follow with food at a reward area each time, before returning the animal to the home cage. Repeat, while increasing block size by one trail each time, until animals reach 10 trials per block.

## 3. Automated feeder habituation

The rat is still rewarded with a pellet for nose-poking, however the nose-poke is now followed by delivery of food at a reward location via the automatic feeders. The rat will need to habituate to the sound of the feeders. Phase out food at the nose poke slowly until the rat is nose-poking and receiving the resulting reward at the reward areas only. Initially a home cage break should be provided after 5 consecutive trials. The amount of trials should be gradually increased to 10 trials. Feeder habituation is complete when the animal has habituated to the sound of the automatic feeders and behaves according to the task structure. It must nose poke to initiate a trial, immediately proceed to the active feeder, consume the reward, and return to the nose poke location to initiate the next trial for at least 10 trials within a session.

## 4. Cue Training

Blocks initially consist of 10 trials per block, a session consists of 4 blocks.

*Phase 1*

1. A tone (randomly selected at each trial from two possibilities) is played in response to nose-poke. Reward the animal with 2 pellets at reward area immediately following the nose poke. Doing this repeatedly allows the rat to become accustomed to the sound and builds the association of sound with reward.
2. The following day, maintain the same trial structure as above, however incorporate a 2 second delay after the nose poke. If the rat reacts by moving towards the correct area reward with 4 pellets. Activate the feeder as soon as the rat makes any motion, however small, towards the correct area. Other behavior results in a reward with one pellet at the correct reward area immediately following the 2 second delay.

*Phase 2*

3. Increase block length to 15 trials.
4. On the first day of phase two reward automatically for nose poke on the first trial, however for half of the remaining trials (randomly chosen) do not automatically reward for nose poke but wait until the rat moves in one direction or the other. Ignore any movement towards the incorrect reward area, and end the trial if the animal turns towards the incorrect area completely. Immediately reward any movement toward the correct reward location with 4 pellets.
5. No longer reward automatically on the second day. Movement towards the correct reward location, minimally a 10 to 30 degrees head-turn should be rewarded with 3 pellets in the cued location. If the animal does not move or moves more than 45 degrees towards the non-cued reward location do not reward and wait until the trial ‘times out’ (5 seconds). The nose-poke will then be available again for the next trial. Move to step 6 when the animal responds to the tone with the desired behavior of a 10 to 30 degree head turn in at least 70% of the trials during a session, for at least 3 sessions.
6. Head movement of minimally 60 degrees results in reward. This response must be present in 70% of the trials within a session before advancing to the next step.
7. More head movement (60 to 90 degrees), and/or some body movement is required (shoulders should turn towards correct reward location). Again, this response must be present in 70% of the trials within a session before advancing to the third phase.

*Phase 3*

8. Rat should be facing correct reward location before reward is given. Advance to phase 4 when this response is present in 70% or more trials within a session.

*Phase 4*

9. Rat should be facing, and once facing have made a move towards the correct reward location before reward is given. Advance to phase 5 when this response is present in 70% or more trials within a session.

## 5. Full Task

A phase is completed when the animal reaches an average of 70% correct.

*Phase 5*

Expand maze, move arms backwards 20 cm so that rats must take the central arm to reach the reward. Increase block length to 20 trials.

*Phase 6*

Move the arms back another 50cm from the startbox and increase the block length to 25 trials.

*Phase 7*

Move the maze arms 80 cm from the startbox.

## Experiment Design

## Technical Implementation

We created an automated training system consisting of the following modules:

- *Input devices* that register the actions of the animal. It can be an infrared (IR), pressure or conductivity sensor. It translates the action into a simple signal, for example a voltage change.
- *Interaction (output) device* such as a food dispenser.
- *The main controller* that enables basic communication between registration and interaction units. It also sends and receives data from the computer running the software.
- *Tracking (input) devices* that register complex data to be analyzed offline on the computer.
- *Software program* that manages the experiment, saves data and provides simple statistics.

Every module has its own unique task and is independent from the other modules. This is important as it reduces the probability of errors occurring and is useful for debugging. Modules can be easily modified and improved without affecting the rest of the system.

The prototype of this system consisted of Arduino Uno microcontrollers functioning as the motherboard and IR sensor board, and Arduino Nano development boards as the microcontrollers for the feeder units. This construction allowed for fast prototyping. A future version of the system could replace the Arduino boards of the slave microcontrollers with custom designed printed circuit boards containing microcontroller chips.

## Hardware and Software Components

### Main Controller

The main controller is an essential element of our system. We used an Arduino Uno as the master in the I2C protocol. Importantly, the experimental trial logic is handled on the main controller. This provides a speed advantage, because communication between the master and “slaves” microcontroller using I2C is much faster that serial port communication with a computer. The Master microcontroller can be accessed by the user via a PC, to make changes during a trial, for example to finish a trial earlier, give clues or give additional reward based on the analysis of the behavior of the animal. I2C extenders provide the ability to connect the master controller to a large number of slave units.

## IR Sensor Board

The sensor board consists of an Arduino Uno microcontroller with IR sensors connected to its digital input pins. When the main controller is waiting for sensor input it requests information regarding sensors triggered ten times per second from the sensor board. Each sensor pair is identified with a number. Once a sensor is triggered the sensor board saves the number assigned to that sensor. This variable cannot be overwritten until the next request from the main controller, during which the number of the activated sensor is sent to the main controller and the variable is reinitialized. This ensures only one sensor reading is received by the main controller per request. When the main controller receives the number of the triggered sensor, it closes the communication with sensor board and the trial is ended.

## Feeder

Each feeder is an independent slave device, driven by microcontroller that receives the number of food pieces to release through I2C protocol. The feeder device consist of a 3D printed pellet reservoir containing a rotating disc, a servo motor and an electronic circuit with IR sensor, I2C extender and microcontroller. The program of the device independently controls the fulfillment of tasks as follows:

1. The feeder microcontroller receives the number of food pellets to dispense.
2. The feeder microcontroller triggers the servo to spin the disk.
3. If no pellet has fallen in certain number of seconds it spins the disk in the opposite direction.
4. The IR sensors in the tube registers the falling pellets
5. If the number of registered pellets reaches order amount, the servo stops.
6. The feeder microcontroller sends information to the master microcontroller (main controller) that food is delivered.

## Software Program

A Python program with a graphical user interface (GUI) is used to run the experiment. The core of this program is the Experiment class, which defines all parameters and functions related to a specific experiment. It is also responsible for storing data. The second layer is GUI interface, which should be intuitive to enable the trainer to intervene in experiment and review the results gathered. The software was developed in Python as this programming language allows for rapid prototyping of code customized for the experiment. The GUI is based on the PyQt5 library.

The Experiment class can also be used without the GUI. It communicates with main controller by serial port and it includes following features:

- A default dictionary of settings (name of experiment, total and per day number of blocks, number of trials in block, number of animals, Arduino serial port address and 3 threshold times discriminating the number of pellets to reward, and an upload settings function, which sends the parameters to master Arduino.
- Save/load experiment settings and data (serialized data structure to write to a file efficiently).
- The randomized stimulus presentation schedule is generated in the form of a list. The randomization can be modified to adjust to specific experiment requirements (for instance, no more than three of the same stimuli presented in a row).
- Running the entire block of trials for given list of randomized locations
- Starting of trial and receiving the result, which is immediately saved to a text file and into the class data container.
- Administering additional reward or canceling the trial at any moment.
- The possibility to add a comment to a trial.

The commands mentioned above can also be issued in the command line if preferred.

## Randomization

The sequence of cues across trials is crucial for successful task learning. A completely random sequence will most likely lead to a negative learning outcome as rats are sensitive to spurious correlations and runs in the trial sequence. Especially in the early stages of learning, rats will respond to perceived patterns in the sequence of cues and reward deliveries. The difficulty is compounded by the fact that in a binary task, a random strategy will lead to reward in 50% of the trials, which may quell motivation for learning the strategy leading to 100% rewards. One of the most common strategies applied by rats in a two-alternative forced-choice task is known as alternation, which involves alternating between the possible reward areas. This strategy may reflect natural foraging behavior where it is not strategic for the animal to return to a depleted food source [2]. The randomization of the trials must be designed to discourage this natural tendency to alternate. It must, however, also not allow the same side to be rewarded too many times in a row since this can result in the development of a bias to the frequently rewarded side. Once such a bias is established it can persist for several sessions and prevent the animal from learning the task. It is therefore crucial to test the randomization algorithm for repetitive patterns and amount of switching between sides, too much of which encourages alternation behavior.

Randomized sequences of stimulus presentation are therefore tested for patterns that may establish a bias either towards one particular side or towards alternation behavior. Sequences are generated at the start of each block. When a sequence does not meet the requirements it is discarded and a new sequence is generated and tested against the requirements.

To examine possible patterns occurring in the generated sequences that meet the requirements the randomization is run many times and the generated sequences analyzed for potential patterns. Here the randomization has been created ten thousand times to observe the frequency in which certain patterns occur within a block of 20 trials.

The number of left and right trials within a block must be balanced. The figure below displays the distributions of the number of trials rewarded on the right side of the maze, and the number those rewarded on left. These distributions must be almost identical to avoid biases.

When many consecutive trials reward a particular side, a bias towards that side may be formed. To avoid such a bias the randomization cannot contain more than three trials in a row of a particular side. However this restraint can result in sequences which contain a relatively high amount of alternation transitions between trials (see figure on the right), encouraging alternation behavior.

Reducing the number of allowed alternation transitions corrects for the natural tendency of the animal to alternate. The randomization simulation resulting from the implementation of this measure does not allow for more than 9 alternation transitions in a block of 20 trials, shown in the figure below.

The amount of left to right versus right to left transitions should also be equivalent.

The number of trials per block varies per phase, therefore these checks should be performed for all block sizes used.

## Strategy Simulations

As mentioned previously rats are excellent in identifying patterns in the randomization in order to predict which side has a high probability of delivering reward in the upcoming trial, given the outcomes of the previous trials [3]. Predictable patterns in the randomization must be avoided as they may be learned by the animal, providing a way to solve the task without attending to the cue, obtaining the reward significantly more than 50% of the times and minimizing effort. Such behavior would hinder the learning of the intended task.

Thus, the use of common strategies (e.g. alternation) must be discouraged by ensuring that these do not result in an excess of obtained rewards. The threshold at which an animal determines that a strategy is viable differs per animal. However as a general rule the randomization should not reward these strategies more than chance level, or 50 percent. Our pilot studies revealed that when adherence to a strategy consistently results in reward over 60 percent of the time animals will be likely to use a strategy. This was especially true for common strategies requiring limited mental resources such as alternation.

Several common behavioral strategies related to patterns in the randomization have been observed in pilot data. Common simple strategies are alternation, or always choosing one particular side. Slightly more complicated are patterns such as left, left, right, or right, right, left, etc. More sophisticated strategies involve using success of a previous decision to determine the next choice of side. For instance an animal that systematically returns to the same location if it was rewarded, and switches to the opposite location when not rewarded is using what is often referred to as a ‘win-stay’ strategy. A ‘win-shift’ strategy involves choosing the opposite location to the one rewarded in the previous trial.

The randomization must not reward these strategies either, therefore it should also be checked against patterns which may facilitate and encourage this behavior. This can be ensured through the simulation of these strategies on the generated randomizations and scoring how often they are successful.

First, simple strategies were simulated, meaning static strategies that are not based on changes in the environment such as the tone, or whether a reward was given at a particular location in a previous trial. Such strategies include consistently choosing only the left or only the right side, alternating between the two sides, or choosing sides based on a set pattern, such as go left twice, then right once, then left twice again, etc. None of these strategies will reward the rat more than 50 percent on average as can be seen in the figure on the right.

A slightly more complex strategy involves responding to a change in the cue tone. In this case the animal switches the chosen reward location when the cue tone has changed from one trial to the next. The use of this strategy indicates that a change in tone between trials is associated with a change of reward location instead of associating a specific tone with a particular side. This strategy can result in a run of successful trials if the animal happens to start on the side correctly associated with the tone. An animal displaying this behavior may appear to have learned the task of associating a particular tone with a specific reward location, when in fact, it has learned a different task, namely to switch to the opposite reward location on a change of tone. On the other hand, an incorrect initial decision in the first trial will result in a block of trials where the animal will not receive any rewards unless a strategy-change occurs within the block of trials. It is also possible for the animal to start out correctly but become distracted during the course of the block and miss a tone change, after which the outcome of the strategy will reverse.

Simulating this switch-on-tone-change strategy, including runs where the simulated rat was distracted (random side was chosen) every 5 or 10 trials, or a random number of trials between 5 and 10 each time, and the tone change strategy was continued, now based on the random response from a ‘distracted’ trial, did not result in success more than 50% of the time on average. Neither did the previously described win-stay and win-shift strategies.

The choices of the animals were analyzed for compliance with common strategies during the course of the experiment. We observed that the natural tendency of the animals towards alternation between reward areas was remarkably strong. A reward percentage of chance level did not deter them from relatively frequently regressing to the use of this strategy, even in later stages of training. Consequently the randomization was adjusted to discourage this tendency in the full task (phases 5 to 7) by increasing the chance that a particular location was rewarded twice in a row to 60 percent, thereby lowering the chance of reward at the opposite location in the next trial to 40 percent. This proved to be enough to break the habitual alternation.

## Animal Scores on Use of Strategy

The simulations of the various strategies discussed earlier indicate that the animals could not use one of these strategies and score above 60 percent. To ensure that the animals did learn the task and were not applying a strategy the choice of the animals for each trial was evaluated based on its choice in the previous trial. That is, the choice of the animal is compared to the choice it would have made if it chose according to a predetermined strategy.

Scoring the reaction of each of the strategy stimulations on the choices made by the animal in the previous trial reveals a similar pattern as was seen in the strategy simulations previously. The figures show the percent of correct choices the animals made during the experiment that would have also been correct if they had used either a win-stay, win-shift, or alternation strategy. Considering the actual scores are much higher we can conclude that animals could not have relied on one of these strategies to achieve these results.

## Training Methods (protocols)

### Animals

Four 6 months old Long Evans male rats were housed in pairs in large cages (610 x 435 x 215mm) maintained on a reversed 24h light/dark cycle, and food deprived to 85% of their ad libitum weight, based on a ad-libitum feeding weight curve by animal supplier Janvier. Training sessions took place during the dark portion of the cycle.

### Food Restriction and Feeding

In order to learn the rats must complete at least 30 trials per day, the rats must be motivated and food deprived to 85% of their ad libitum weight.

Food intake was restricted to 15 grams per day for each rat, of which on average 10 grams were earned as reward in the maze during the task, and the remaining food (15 grams minus reward earned in maze) fed in the home cage. In group housed animals however, this may be problematic, as the dominant rat tends to eat more, decreasing his motivation the next day, while the non-dominant animal will not have a chance to eat the amount of food he needs each day. To prevent this, food is spread over the cage (for instance a few pellets in every corner) so that while the dominant animal is eating at one end of the cage, the other animal can eat elsewhere in the cage out of reach of the dominant animal. Alternatively, animals may be fed separately, however this requires habituating them to separate cages for feeding. For optimal results more blocks of trials should be added per training day, with rest periods between blocks so that the entire daily need of 15 grams is provided in the maze via the task as reward.

Pellets fed as rewards were 45mg Supreme Mini-Treats from Bio-Serv. In this particular experiment a mix of bacon and apple flavors was used, however any flavor, or non-flavored pellets also suffice[4]. Larger pellets are not recommended however as they decrease the amount of trials that can be run daily.

Data were taken from average weight per week of age provided by the supplier Janvier and supplemented by our own data from eight animals from Janvier previously used in experiments for weeks 25 to 40 as Janvier could not provide weights for this age range. Animals were weighed three times per week. These weights were averaged and plotted per animal on this growth curve chart weekly. Weights should fall within the purple shaded error band around the 85% curve. When this was not the case the home cage feeding was adjusted accordingly.

### Task Design

Trials are grouped in blocks, the number of trials per block varies per phase. A rat remains in the maze for the duration of a block and is then placed back in the home cage. Meanwhile the maze is cleaned, and another rat is placed in the maze. This block design allows animals to rest, drink, and regain hunger and motivation in between blocks. Training with 4 animals per group creates a sufficiently long recovery time for the animals to regain motivation before entering the maze again for the next block. In this experiment the animals had four blocks of trials per day. Initially the animals may still be wary of the new environment, thus the blocks in the first phase should consist of 10 trials only in order to provide sufficient pauses in the training. As the animals become more at ease in the environment and accustomed to the task and the daily routine the amount of trials increases to 15 per block in phase 2, then 20 trials per block, and finally 25 trials per block.

When pair housed often one animal will be dominant over the other. This dominance is often displayed in the form of the dominant animal pinning down its cage mate on its back. It is best to always take the dominant animal first as natural rat behavior dictates that the dominant animal, because in natural situation that animal would have priority in exploration and access to food [5]. If the non-dominant animal is placed in the maze before the dominant animal, the dominant animal may act aggressively upon his return to the home cage.

In experimental designs that do not require counterbalancing for this aspect, placing animals in the maze in the same order everyday will help them establish a routine, which may contribute to reduce stress.

### Monitoring performance

Performance was monitored daily. At the end of a training day the performance of each rat was plotted and compared to previous days. In this task thresholds to obtain reward are altered incrementally. As the requirements for obtaining a reward becoming increasingly demanding it is to be expected that performance may decrease somewhat with a change in requirements. For some animals however the increased difficulty may result in a drop in obtained reward, and engagement in the task. Because of this, the previously required behavior was rewarded, though with fewer pellets. For instance for the previously desired behavior reward two pellets, and for the newly desired behavior reward four. We found that even an increase by one pellet for ‘better’ behavior will quite consistently positively bias the rats towards displaying that behavior more often. Dominant animals may, in response to receiving less pellets, choose to apply an easy strategy such as alternation [6], or exhibit decreased motivation. In those cases, we avoided feeding animals in the home cage where one animal may eat more than the other, and all food was provided on the maze, to each animal separately. Alternatively, the threshold may be lowered temporarily to such a degree that the rat is capable of achieving it, to then be slowly be increased in difficulty again.

The reaction time was also used to determine the amount of reward. The standard reward for a correct choice within the time limit was two 45 mg pellets. One extra pellet was given for reaction times within three seconds. This time limit was based on observation. Longer reaction times often indicated hesitant behavior or lack of attention. In this manner, purposeful and goal directed behavior was encouraged. In the first two stages an extra pellet was also rewarded if the animals exceeded their criteria for reward. For instance, if the criteria for a reward was a 30-degree turn, then a 60 degree turn was rewarded with an extra pellet.

The first and second phase of the cue training included ‘hint’ trials where one reward was given immediately following the cue tone. On the first day of the first phase all trials consisted of hint trials. The number of hint trials was gradually decreased over several days. Near the end of phase two hint trials were rarely used. After these first two phases hints were only given occasionally to improve motivation. These hint trials excluded in the manual and reward based scores.

## Analysis Methods

The goal of this study was to develop and test a fully automated system, including scoring of trials. To test the accuracy of automatic scoring, all trials were scored manually as well as automatically. Manually each trial was scored as correct, incorrect, or canceled. A trial was invalid or canceled if the animal did not react to the cue within the time limit or the animal did not leave the start-box before the time limit was reached, in case of hardware or software malfunction, for instance if the cue was not given, or due to human error (If, for example, the researcher interfered with the trial accidently), or if a hint was provided in the form of one pellet dispensed from the reward area immediately following the tone. This was necessary at times to calm an anxious animal, or to motivate an unmotivated animal. For number of valid trials per day see figure 1 in the supplementary materials.

For each trial the training program records which side was cued, which reward area sensors the rat activated first, the time between the activation of the nose poke sensors and the activation of the reward area sensors (reaction time), how many pellets were rewarded by the computer, and how many extra pellets were rewarded manually by the operator.

Several different scoring methods were tested and compared. The first consisting of summing the trials marked manually as correct. Also, the trial information gathered by the software was used to compare the cue side with the reward area sensors that were activated first. If these were identical then the trial was marked as correct, if they differed the trial was marked as incorrect, and if the trial was marked as not valid then neither label was assigned. The time limit within which a trial was valid varied per phase.

The compact size of the maze during the first phases presented a problem as it was not uncommon for the tail of the rat to activate the sensors. Consequently, sensor information was not always reliable in the first four phases. To score the automatically-gathered trial data correctly without relying on the sensor activation required examining the rewards given. In phase 1, the criteria for a correct trial was at three or more rewarded pellets. This served not only to motivate the animals but also for feeding purposes, to ensure they received the majority of their daily food intake in the maze, as the first phase consisted of only 30 trials per day. In phase 2 to 5, two or more pellets, and in phase 6 and 7 one or more pellets.

Not all training days contained an equal amount of valid trails per day. Trials could be declared invalid for reasons such as the occurrence of a technical error, or the inadvertent activation of the nose-poke sensors. Supplementary figure 11 provides an overview of the amount of valid trials per day for each animal (average over animals in black).

## Supplementary References

**Supplementary Table 1:** Settings for each phase defining the length of the central arm in centimeters, number of trials per day, time allowed (in seconds) for a trial, minimum number of pellets to be rewarded for a correct answer, and the timing thresholds number of extra pellets dispensed for response windows rewarding faster reaction times.

|  |  | Central Arm Length (cm) | Trials Per Day | Time Limit (s) | Min. Reward (num. pellets) | R+ time limit (s) | R+ (num. pellets) | R++ time limit (s) | R++ (num. pellets) |
| --- | --- | --- | --- | --- | --- | --- | --- | --- | --- |
| Cue Training | Phase 1 | 0 | 30 | 6 | 2 | 5 | 4 | 3 | 5 |
|  | Phase 2 | 5 | 60 | 6 | 1 | 5 | 2 | 3 | 3 |
|  | Phase 3 | 5 | 60 | 5 | 1 | 4 | 2 | 3 | 3 |
|  | Phase 4 | 5 | 60 | 5 | 1 | 4 | 2 | 3 | 3 |
| Full Task | Phase 5 | 20 | 60 | 6 | 1 | 4 | 2 | 3 | 3 |
|  | Phase 6 | 50 | 80 | 6 | 1 | 4 | 2 | 3 | 3 |
|  | Phase 7 | 80 | 100 | 6 | 1 | 4 | 2 | 3 | 3 |

**Supplementary Figure 1.**
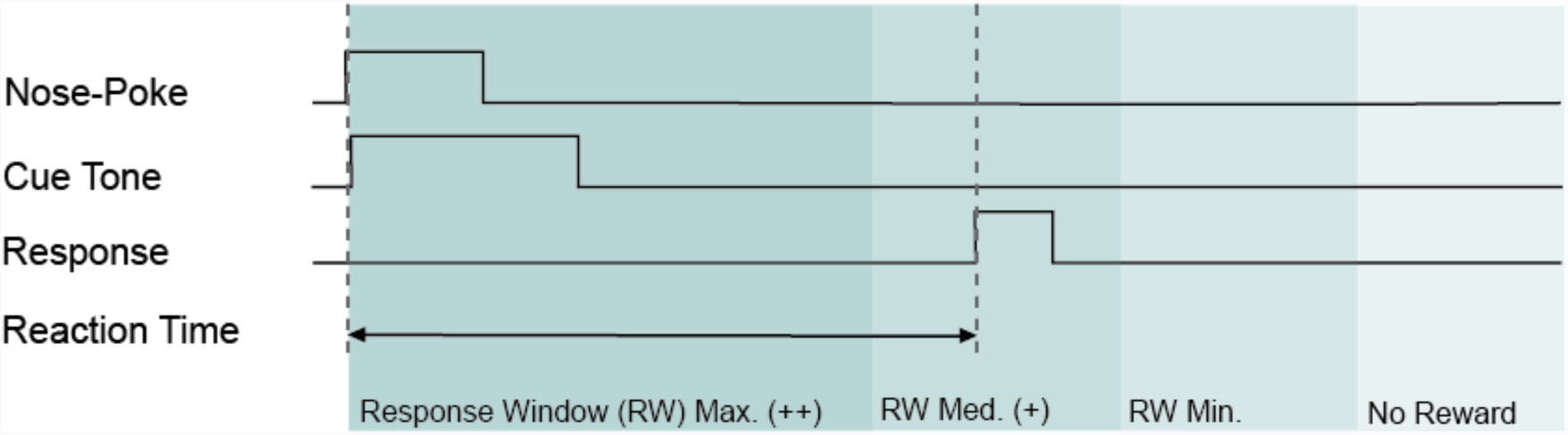
Trial structure for cue training (response window depend reward from phase 3 onwards) and the full task. A trial begins with a nose poke, which triggers the playing of a cue tone for 1 second. The reaction time is measured from the onset of the cue tone to the activation of the sensors at the reward location. Several response windows are defined that determine the number of pellets dispensed, encouraging the animals to respond to the cue in a timely manner. Faster responses are rewarded more than slower responses. If the animal responds slower than the time allowed no reward is given.

**Supplementary Figure 2.**
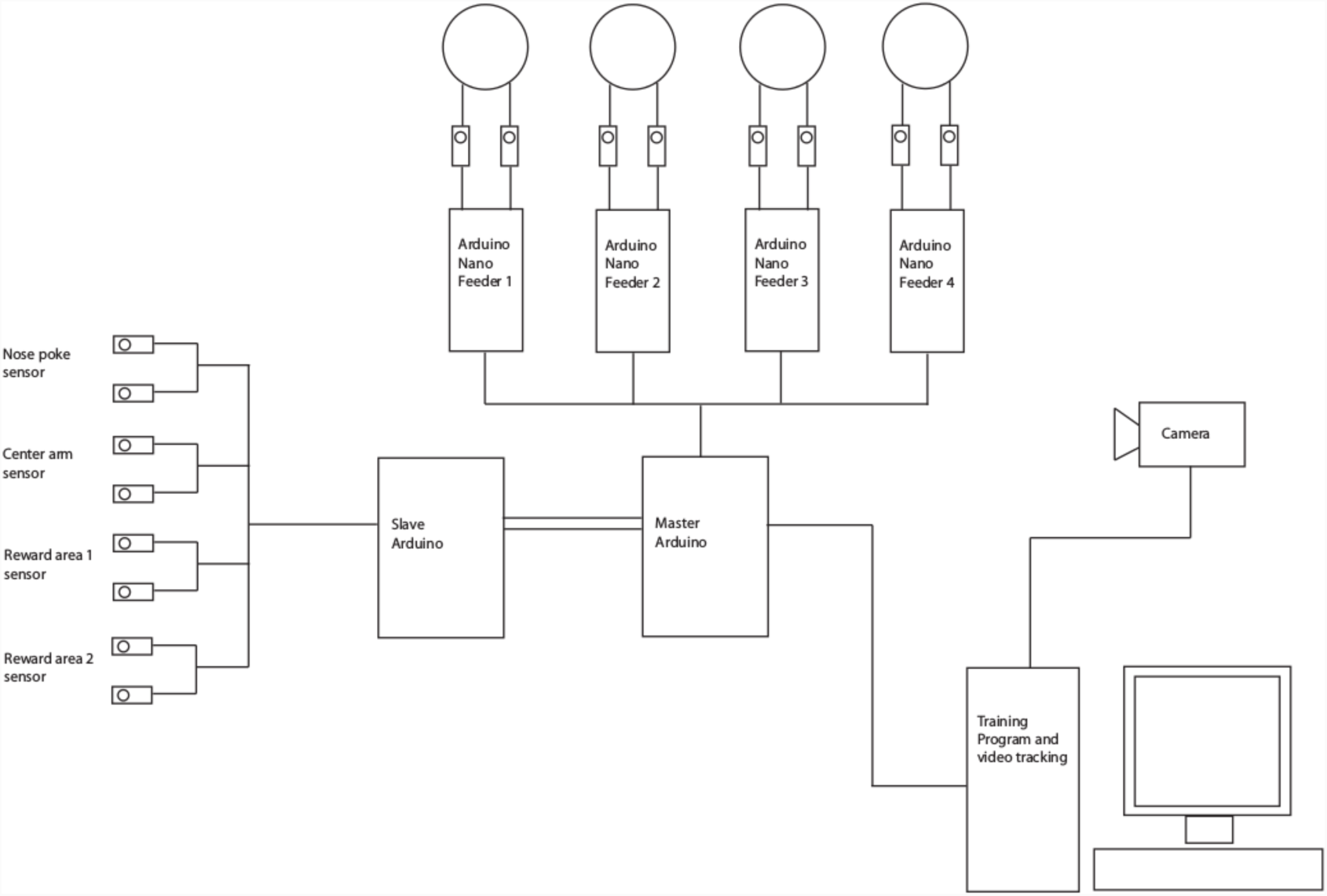
Hardware schema showing the computer running the training software and video tracking, the camera recording the trials, the main microcontroller acting as a master and connected to the four feeder units, each controlled and monitored by their own microcontroller, and also connected to another microcontroller that registers the input from the four infrared beam sensors pairs.

**Supplementary Figure 3.**
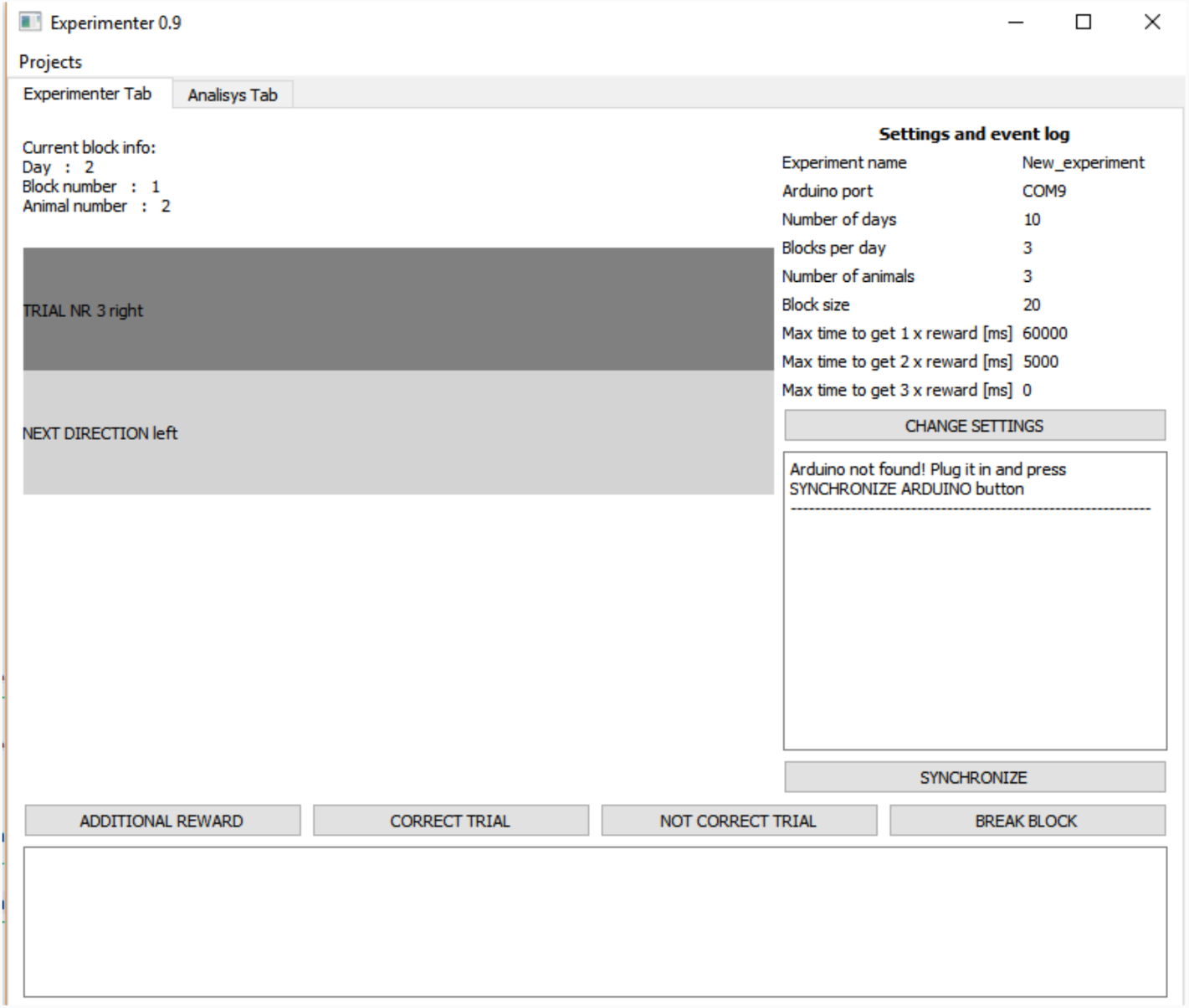
Screenshot of the software program showing the information regarding trail in progress including the training day, the block number, and the number identifying the animal. The current trial is shown underneath this information in a rectangle colored according to the rewarded side. A preview of the next trial is shown in the rectangle underneath. The experiment settings are shown in the left hand column including the name of the experiment, the name of the connected microcontroller, the blocks per day, the number of animals, the number of trials per block, and the maximum time allowed defining the three response windows. The ‘change settings’ button directly underneath allows the user to adjust these settings. The window below displays feedback received from the microcontroller and the button below this window allows the user to manually synchronize the microcontroller in case the automatic synchronization experienced problems. The four buttons spanning across the screen horizontally facilitate manual control of the maze including the ability to dispense extra pellets, mark a trial as correct or incorrect, and end the block, in case the animal has lost interest and is no longer initiating trials. In the area below the user can write notes regarding any unusual events or comments regarding the trial if necessary.

**Supplementary Figure 4.**
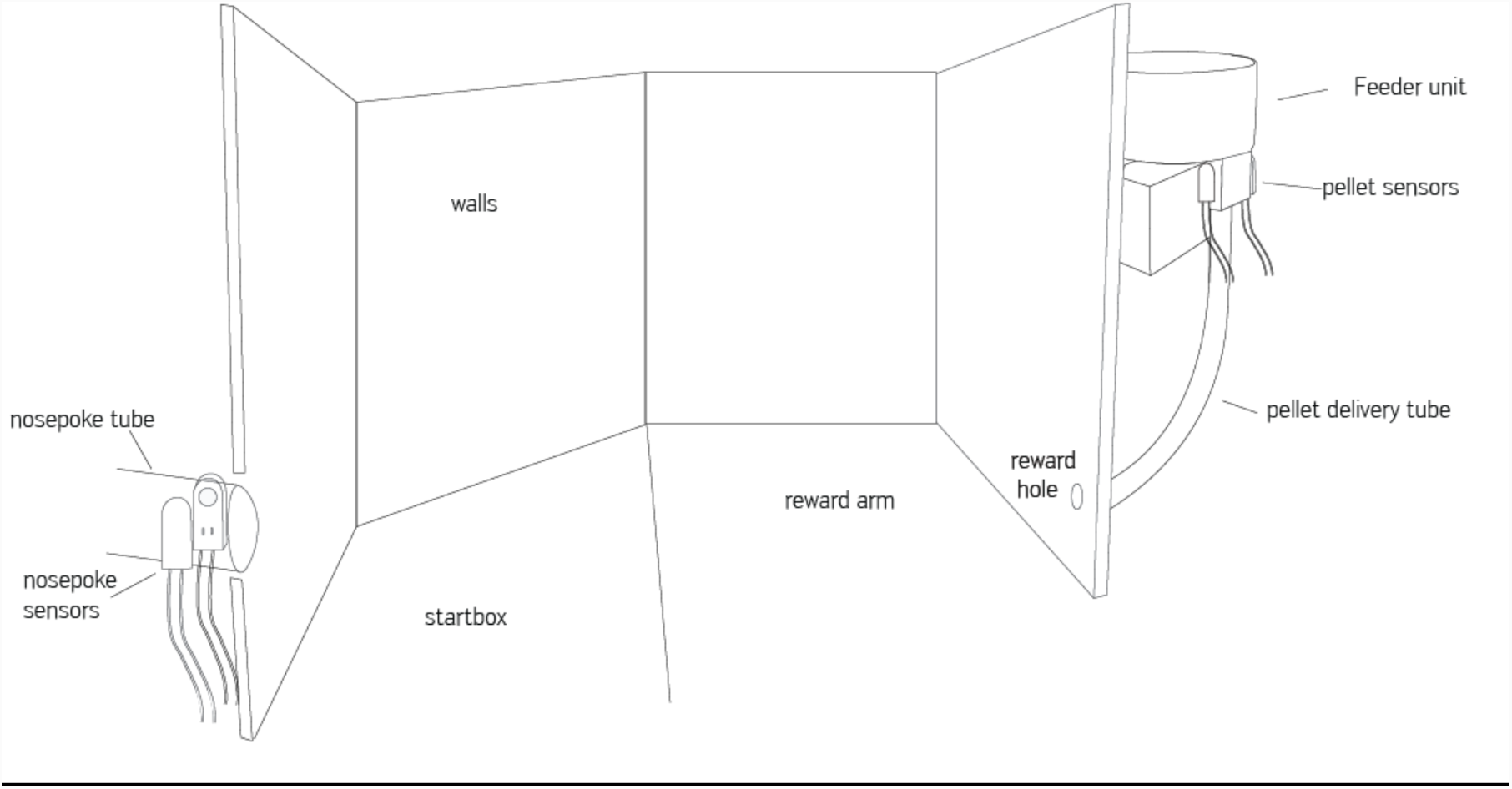
The maze set up at the first phase including 40 cm high walls, nose poke tube with infrared beam sensors. The reward arm positioned immediately against the startbox and the feeder unit behind the wall consisting of a pellet basin connected to a servo motor and a pellet delivery tube, which delivers the pellets to the animal in the maze.

**Supplementary Figure 5.**
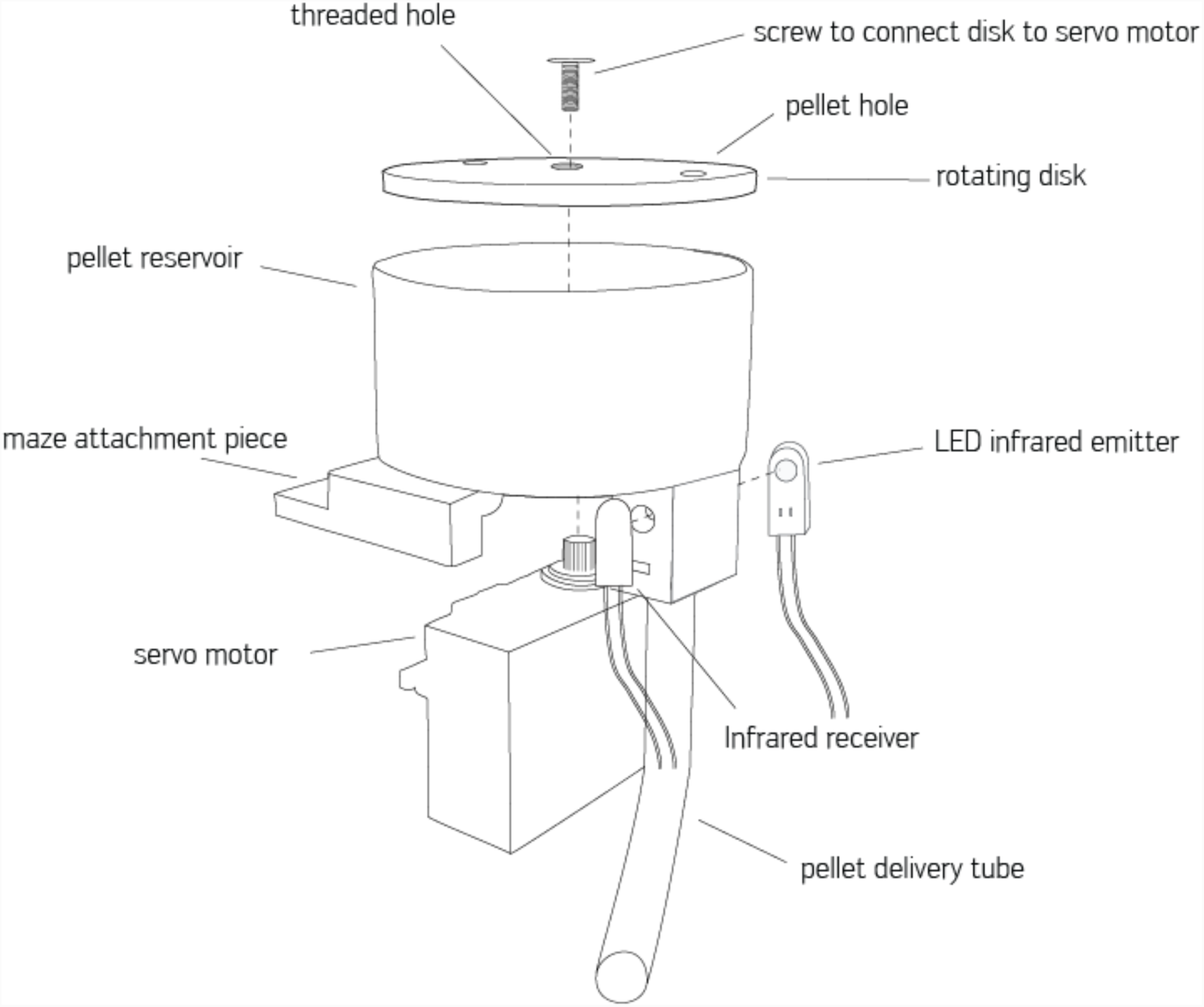
Feeder unit parts include a rotating disk containing two pellet holes connected to the servo motor with a screw. The bottom of the pellet reservoir contains one hole through which a pellet falls when a pellet hole on the disk aligns with it. An extruding piece of the reservoir containing two holes for screws allows the feeder unit to be attached to the maze. The infrared beam created between the LED infrared emitter and receiver detects pellets dispensed into the delivery tube. The microcontroller attached to the feeder unit receives the number of pellets to be dispensed from the main microcontroller and spins the disk until the required number have pellets have been detected by the sensors.

**Supplementary Figure 6.**
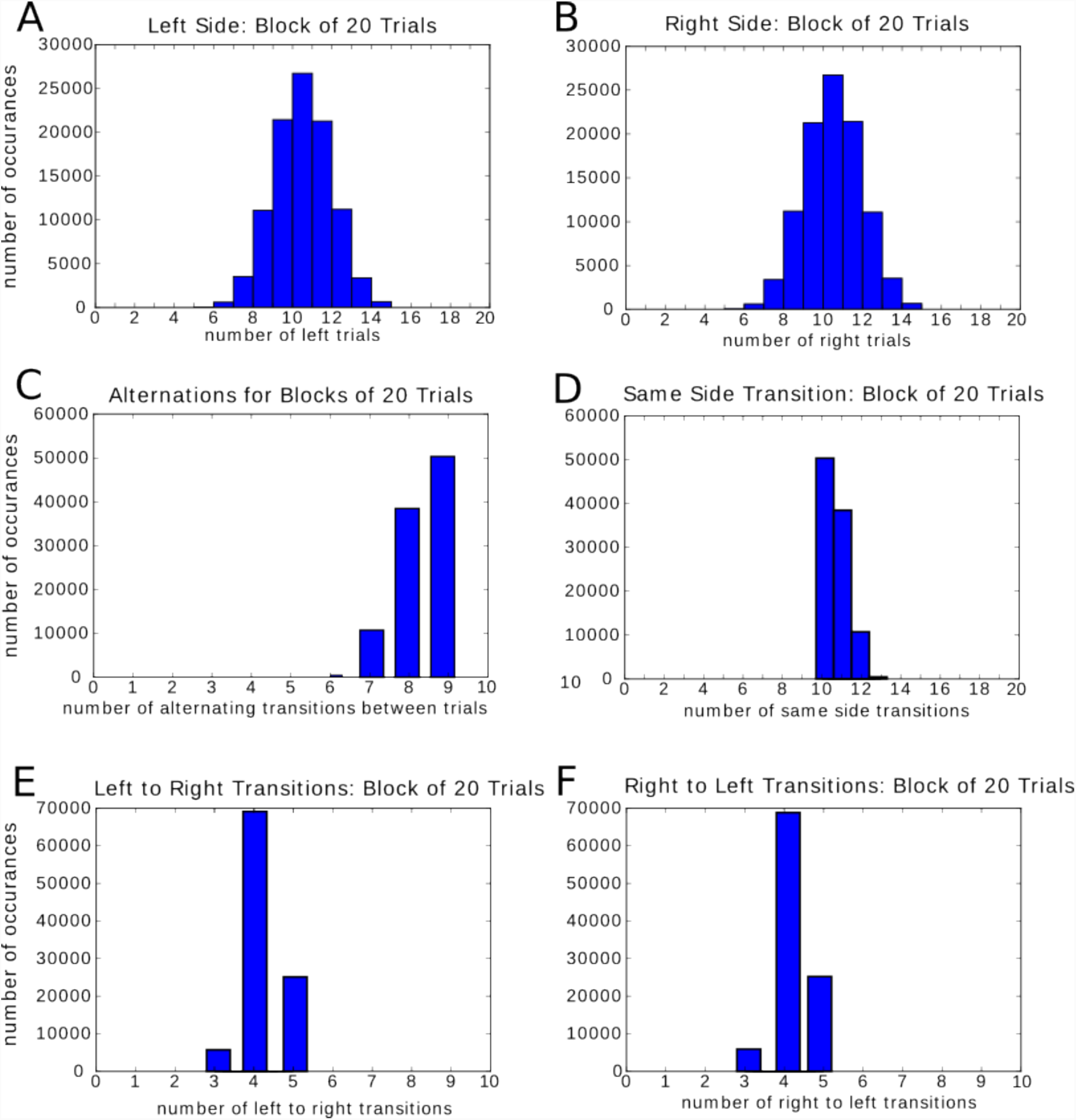
Testing the randomization for patterns. Randomization for blocks of 20 trials simulated 100,000 times. The number of simulated occurrences (y-axis) are plotted for the number of trials where the phenomenon mentioned in the title (for instance: ‘trial is left’, or ‘trial side is identical to previous side in previous trial’ (same side transition)) occurring in each simulated block. The distributions of the amount of left and right trials occurring in a block are identical to prevent any biases from developing. With two possible choices the transition from one trial to the next can either be to the same side or to the opposite side. The likelihood of the next trial being to the opposite side should be slightly lower than the chance that it is to the same side in order to prevent the animals from alternating between locations on every trial, a common bias seen in rats. Distributions of left to right and right to left transitions should be identical in order to prevent bias forming.

**Supplementary Figure 7.**
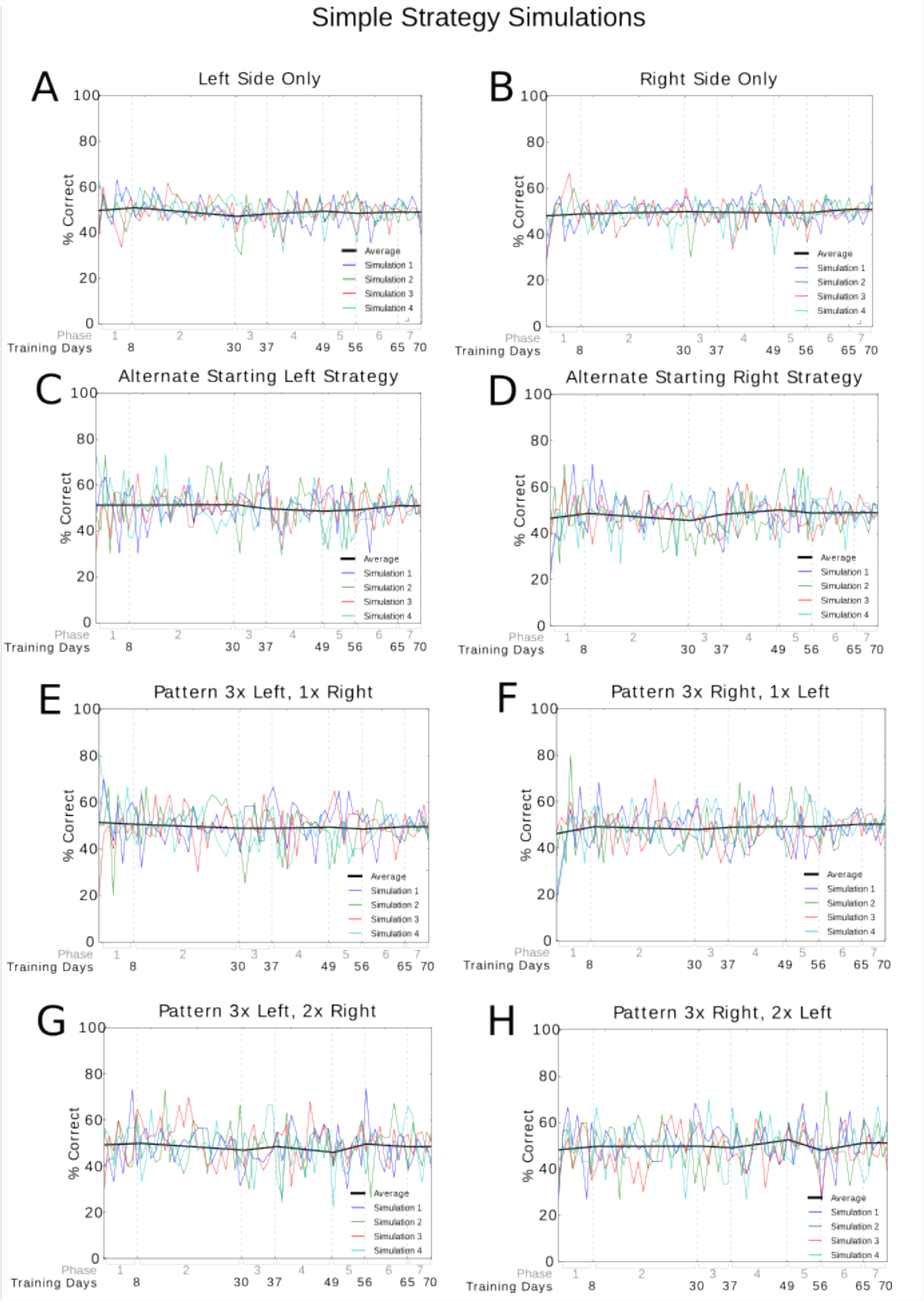
Simple strategy simulations. Simulated rats choosing sides according to a predetermined strategy are presented with sequences of cues generated by the same randomization sequence used in the experiment. Percentage of correct choices (y-axis) are plotted for each training day (x-axis). None of the simulated strategies result in a score higher than 55% on average.

**Supplementary Figure 8.**
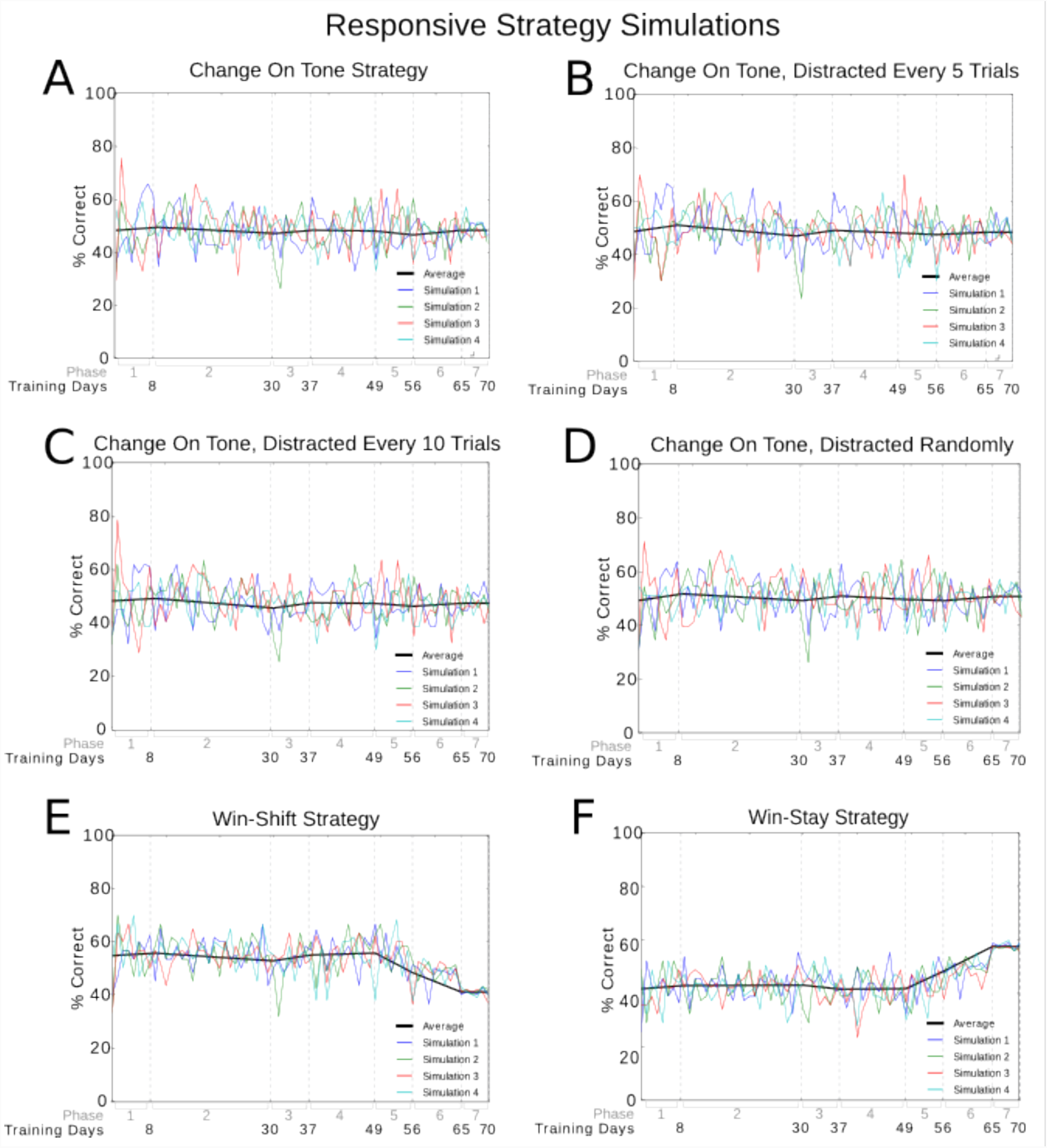
Responsive strategy simulations. Percentage of correct choices (y-axis) are plotted for each training day (x-axis). Simulations responded to sequences of cues, generated with the same algorithm used in the experiment, according to a particular strategy. The simulations were programmed to change their choice in side in response to a change in the cue tone (with several variations such as ‘distractions’ occurring every 5 or 10 trials, or a random number between 5 and 10. The win-shift and win-stay strategy simulations were programmed to change sides related to their success in the previous trial. In win-shift that means if the choice the simulation made in the previous trial was correct then the opposite side was chosen in the next trial, and an incorrect trial resulted in a choice for the same side as chosen in the previous trial. For win-stay the same side was chosen again following a correct trial, and the opposite following an incorrect trial. These two strategies disregard the cue tone altogether. None of the above strategies resulted in a score higher than 60%.

**Supplementary Figure 9.**
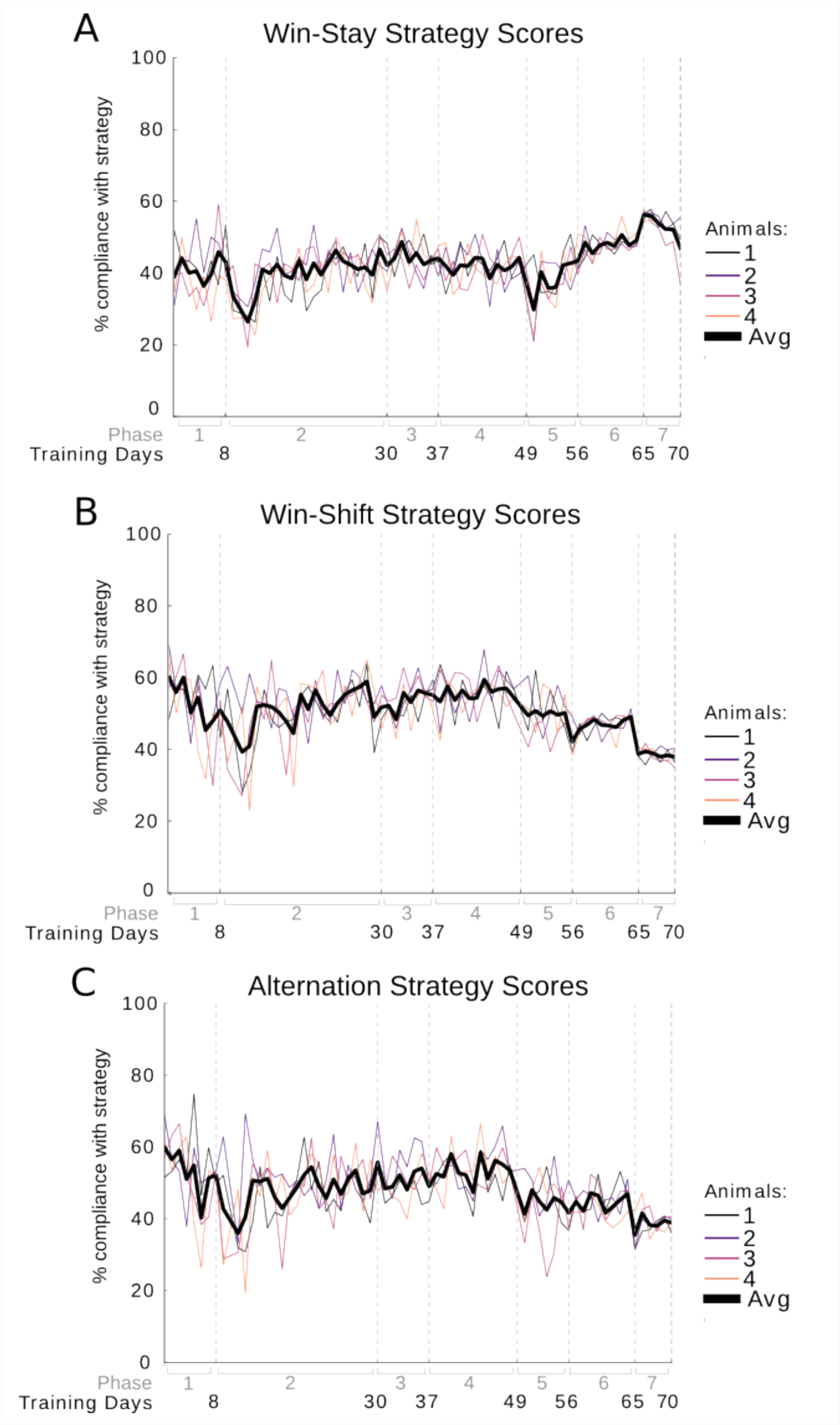
Percent compliance of animal choices with win-stay, win-shift, and alternation strategies. Percent compliance is shown on the y-axis and the training days and training phase on the x-axis. Data for each animal is plotted separately and per day as well as the average over all animals per day (thick black line).

**Supplementary Figure 10.**
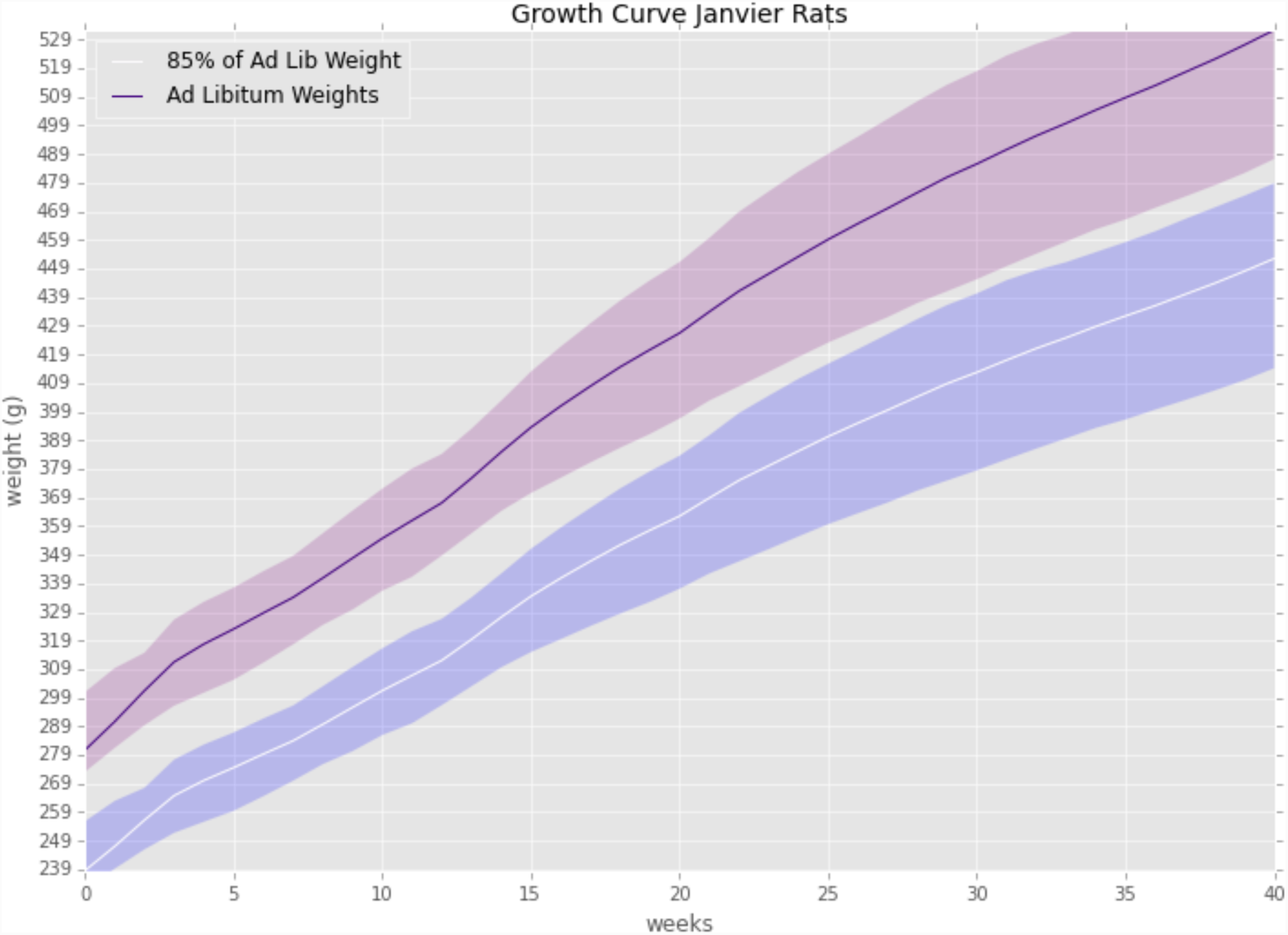
Growth curve showing the weight in grams on the y-axis and the age in weeks on the x -axis. Data were taken from average weight per week of age provided by the supplier Janvier and supplemented by our own data from eight animals from Janvier previously used in experiments for weeks 25 to 40 as Janvier could not provide weights for this age range. Animals were weighed three times per week. These weights were averaged and plotted per animal on this growth curve chart weekly. Weights should fall within the purple shaded error band around the 85% curve. When this was not the case the home cage feeding was adjusted accordingly.

**Supplementary Figure 11.**
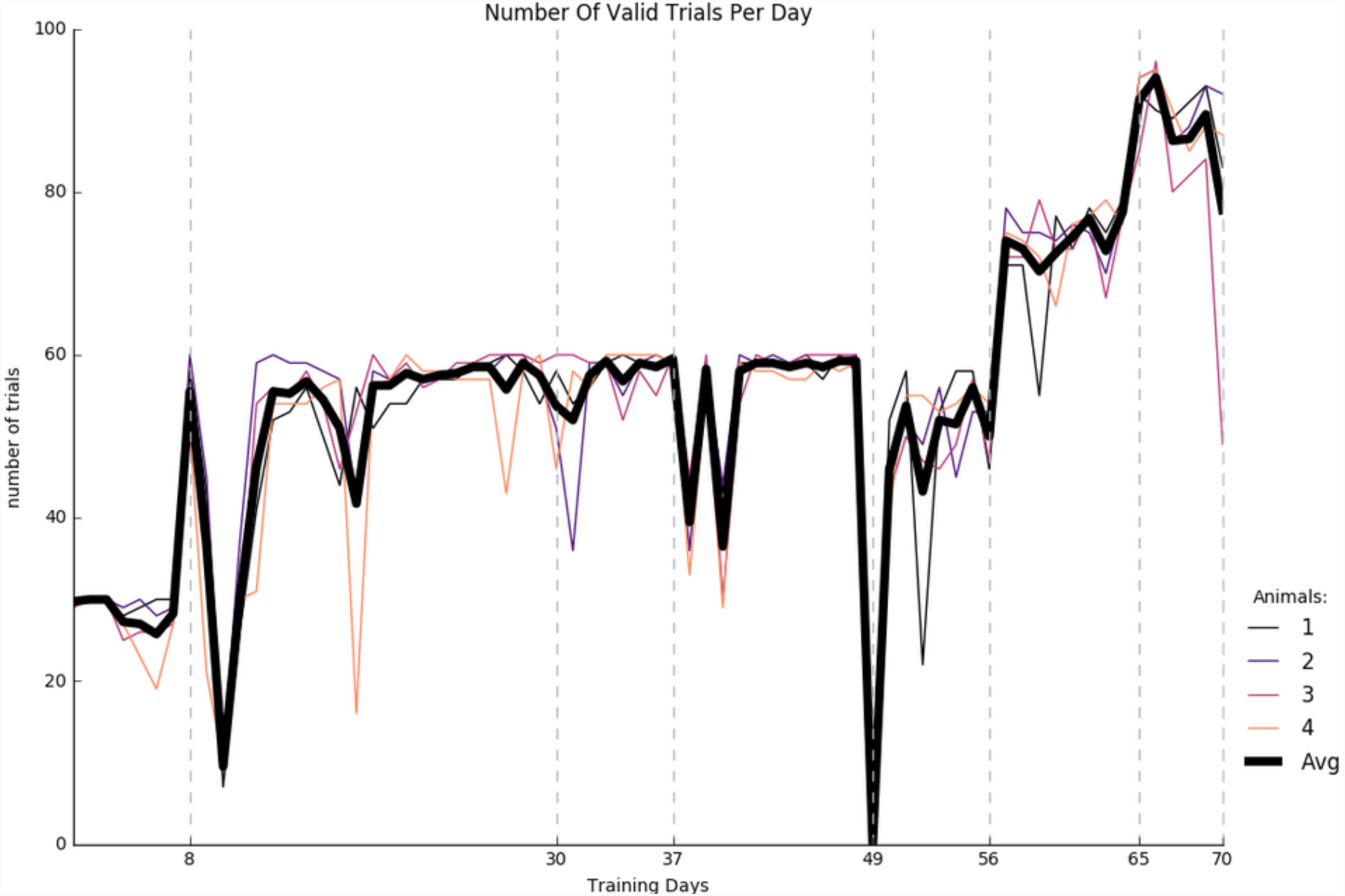
Number of valid trials (y-axis) per day (x-axis). The large dip at day 49 was due to a hardware failure due to which no training could take place that day. The Python and Arduino code necessary to run an experiment using the automated training method described in this paper can be found at: https://github.com/estherholleman/AutoTrainProgram

## References

1. Dudchenko P. A. An overview of the tasks used to test working memory in rodents. Neurosci Biobehav Rev 28, 699–709 (2004).

2. Vorhees C. V., Williams M.T. Assessing spatial learning and memory in rodents. ILAR J. 55, 310–32 (2014).

3. Morris, R. Developments of a water-maze procedure for studying spatial learning in the rat. Journal of Neuroscience Methods, 11, 47–60 (1984).

4. O’Keefe J, Dostrovsky J. The hippocampus as a spatial map. Preliminary evidence from unit activity in the freely-moving rat, Brain Res 34, 171–5 (1971).

5. Bower, M. R., Euston D. R., McNaughton B. L. Sequential-context-dependent hippocampal activity is not necessary to learn sequences with repeated elements. Journal of Neuroscience 25, 1313–23 (2005).

6. Fujisawa, S., Amarasingham, A., Harrison, M. T., & Buzsaki, G.. Behavior-dependent short-term assembly dynamics in the medial prefrontal cortex. Nat Neurosci 11, 823–833, https://doi.org/10.1038/nn.2134 (2008).

7. Wood, E. R., Dudchenko, P. A., Robitsek, R. J., & Eichenbaum, H. Hippocampal neurons encode information about different types of memory episodes occurring in the same location. Neuron 27, 623–633 (2000).

8. Rosenthal, R. Experimenter Effects in Behavioral Research. (Appleton-Century-Crofts, 1966).

9. Pfungst O. Das Pferd des Herrn von Osten (Der Kluge Hans). Ein Beitrag zur experimentellen Tier-und Menschen-Psychologie. (Johann Ambrosius Barth, 1907)

10. Skinner B. F. The Behavior of Organisms: An Experimental Analysis. (Appleton & Company, 1938).

11. Bussey T, Padain T. L., Skillings E. A., Winters B. D., Morton A. J., Saksida L. M. The touchscreen cognitive testing method for rodents: how to get the best out of your rat. Learn Mem 15, 516–523 (2008).

12. Zheng W., Ycu E. A. A fully automated and highly versatile system for testing multi-cognitive functions and recording neuronal activities in rodents”. J Vis Exp 63, https://doi.org/10.3791/3685 (2012).

13. Poddar R., Kawai R., Ölveczky B. P. A Fully Automated High-Throughput Training System for Rodents. PLOS ONE 8 https://doi.org/10.1371/journal.pone.0083171 (2013)

14. Pioli E. Y. An automated maze task for assessing hippocampus sensitive memory in mice. Behavioural Brain Research 261, 249–257 (2014).

15. Holmes A., Rodgers R. J. Influence of spatial and temporal manipulations on the anxiolytic efficiacy of chlordiazepoxide in mice previously exposed to the elevated plus-maze. Neuroscience & Biobehavioral Reviews 23, 971-980 (1999)

16. Balcome, J. P., Barnard N. D., Sandusky, C. Laboratory routines cause animal stress. Journal of the American Association for Laboratory Animal Science 43, 42-51 (2004)

17. Dember W. N., Fowler H. Spontaneous alternation behavior. Psychological Bulletin 55, 412– 428 (1958).

18. Sanderson D. J., Bannerman D. M. The role of habituation in hippocampus-dependent spatial working memory tasks: evidence from GluA1 AMPA receptor subunit knockout mice. Hippocampus 22, 981–94 (2012).

19. Morellini F. Spatial memory tasks in rodents: what do they model? Cell Tissue Res 354, https://doi.org/10.1007/s00441-013-1668-9 (2013)

20. Fam J., Westbrook F., Arabzadeh E. Dynamics of pre-and post-choice behaviour: rats approximate optimal strategy in a discrete-trial decision task. Proc R Soc B 282, https://doi.org/10.1098/rspb.2014.2963 (2015).

21. Schmidt, B., Wikenheiser, A. M., & Redish, A. D.. Goal-Directed Sequences in the Hippocampus. In Goal-Directed Decision Making (ed. Morris, R.,Bornstein, A., Shenhav, A.) 125-151 (Elsevier, 2018).

22. Heredia-López, F. J., Álvarez-Cervera, F. J., Collí-Alfaro, J. G., Bata-García, J. L., Arankowsky-Sandoval, G., & Góngora-Alfaro, J. L. J. B. r. m.. An automated Y-maze based on a reduced instruction set computer (RISC) microcontroller for the assessment of continuous spontaneous alternation in rats. Behav Res Methods 48, 1631–1643 (2016).

23. O’Leary, J. D., O’Leary, O. F., Cryan, J. F., & Nolan, Y. M. A low-cost touchscreen operant chamber using a Raspberry Pi™. Behav Res Methods 50, 1–8 (2018).

24. Zhang, Q., Kobayashi, Y., Goto, H., & Itohara, S. An Automated T-maze Based Apparatus and Protocol for Analyzing Delay-and Effort-based Decision Making in Free Moving Rodents. J Vis Exp 138, https://doi.org/10.3791/57895 (2018).

25. Euston, D. R., Tatsuno, M., & McNaughton, B. L.. Fast-forward playback of recent memory sequences in prefrontal cortex during sleep. Science, 318, 1147–1150 (2007).

26. Wilson, J. W., The phi-maze: A versatile automated T-maze for learning and memory experiments in the rat. Behavior Research Methods, Instruments, & Computers 28, 360–364, (1996).

27. Barnett, S. A. The Rat: A Study in Behavior. (Aldine Transaction Publishers, 1963).

28. Lester, D. Exploratory behavior of dominant and submissive rats. Psychon Sci 9: 285. https://doi.org/10.3758/BF03332224 (1967).

## Supplementary References

1. Modlinsk K. Stryjek R., Pisula W. Food neophobia in wild and laboratory rats (multi-strain comparison). Behavioural Processes 113, 41–50 (2015).

2. Sanderson D. J. Bannerman D. M. The role of habituation in hippocampus-dependent spatial working memory tasks: evidence from GluA1 AMPA receptor subunit knockout mice. Hippocampus 22, 981–94 (2012).

3. Fam J., Westbrook F. Arabzadeh E. Dynamics of pre-and post-choice behaviour: rats approximate optimal strategy in a discrete-trial decision task. Proc R Soc B 282, https://doi.org/10.1098/rspb.2014.2963 (2015).

4. Powell N. J., Redish A. D. Representational changes of latent strategies in rat medial prefrontal cortex precede changes in behavior. Nature Communications 7, https://doi.org/10.1038/ncomms12830 (2016).

5. Barnett, S. A. The Rat: A Study in Behavior. (Aldine Transaction Publishers, 1963).

6. Lester, D. Exploratory behavior of dominant and submissive rats. Psychon Sci 9: 285. https://doi.org/10.3758/BF03332224 (1967).

